# Computational stabilization of a non-heme iron enzyme enables efficient evolution of new function

**DOI:** 10.1101/2024.04.18.590141

**Authors:** Brianne R. King, Kiera H. Sumida, Jessica L. Caruso, David Baker, Jesse G. Zalatan

## Abstract

Directed evolution has emerged as a powerful tool for engineering new biocatalysts. However, introducing new catalytic residues can be destabilizing, and it is generally beneficial to start with a stable enzyme parent. Here we show that the deep learning-based tool ProteinMPNN can be used to redesign Fe(II)/αKG superfamily enzymes for greater stability, solubility, and expression while retaining both native activity and industrially-relevant non-native functions. For the Fe(II)/αKG enzyme tP4H, we performed site-saturation mutagenesis with both the wild-type and stabilized design variant and screened for activity increases in a non-native C-H hydroxylation reaction. We observed substantially larger increases in non-native activity for variants obtained from the stabilized scaffold compared to those from the wild-type enzyme. ProteinMPNN is user-friendly and widely-accessible, and straightforward structural criteria were sufficient to obtain stabilized, catalytically-functional variants of the Fe(II)/αKG enzymes tP4H and GriE. Our work suggests that stabilization by computational sequence redesign could be routinely implemented as a first step in directed evolution campaigns for novel biocatalysts.

## Introduction

Directed evolution is a powerful method to generate enzymes for new chemical transformations.^1,2^ However, catalytic functional groups often have destabilizing effects on protein structure, and altering active site groups for new reactions can lead to unstable, non-functional proteins.^3–11^ Initiating a directed evolution campaign from a stabilized variant can be an effective way to overcome this problem.^12,13^ Typically, a starting point for directed evolution is obtained by screening a library of candidate enzymes for a desired promiscuous activity. If a thermostable homolog with similar catalytic properties is identified, it can then be used as a starting point for evolution of the desired function.^14,15^ Alternatively, there are a variety of strategies to produce stable variants using directed evolution,^16,17^ ancestral reconstruction,^14^ protein recombination,^18,19^ or computational engineering.^20–22^ These methods are often time- and resource-intensive, highlighting the need for simple and accessible alternatives.

An important class of enzymes where stabilization would be useful is the non-heme iron(II) α-ketoglutarate-dependent oxygenase (Fe(II)/αKG) superfamily. These enzymes have emerged as a rich source of potential new biocatalysts. Fe(II)/αKG enzymes can perform asymmetric C-H oxyfunctionalization reactions on small molecule substrates, which could allow expedient and sustainable diversification of simple building blocks to a range of complex polyfunctional compounds.^23–26^ The advantages offered by this enzyme family include a high degree of chemical flexibility in the iron-containing active site due to multiple open coordination sites, the utilization of benign molecular oxygen as an oxidant, and use of the inexpensive and readily available co-factor αKG. However, Fe(II)/αKGs can be relatively unstable,^27,28^ which may limit their practical applications in organic synthesis.

The recent use of an Fe(II)/αKG in an industrial-scale drug biosynthesis pathway highlights both the potential advantages and drawbacks of this family for biocatalysis. An engineered Fe(II)/αKG was used to catalyze an enantioselective C-H hydroxylation to produce a key intermediate for the anti-cancer drug belzutifan.^28^ The reaction could be performed on kilogram-scale and bypassed five steps of the pre-existing chemical synthesis route. Notably, this effort required an extensive, large-scale directed evolution campaign. Furthermore, early rounds of screening yielded stabilizing mutations before significant improvements in turnover could be obtained in later rounds. These findings highlight the potential utility of Fe(II)/αKG for practical, industrial-scale green chemistry but also the importance of enzyme stability in the evolution of new function.

Recent applications of deep learning to protein design have provided new and relatively straightforward methods to stabilize protein scaffolds,^22,29,30^ and there is broad interest in applying these approaches to directed evolution.^31^ Here we demonstrate that the deep learning-based tool ProteinMPNN^29,30^ enables more efficient optimization of a synthetically-relevant, non-native C-H hydroxylation reaction in an Fe(II)/αKG family member. A critical step was restricting the redesign from modifying active site and adjacent residues, which would otherwise be readily mutated to stabilize the enzyme. With a stabilized starting point for site-saturation mutagenesis, we observed substantially larger increases in non-native activity compared to the same mutations in the wild-type parent enzyme. We suggest that this designed stabilization approach should be routinely used in future directed evolution campaigns with the Fe(II)/αKG superfamily and will likely be effective in a broad range of other enzyme families.

## Results and Discussion

### Fe(II)/αKGs with promiscuous activity for C-H hydroxylation of free carboxylate substrates

Within the Fe(II)/αKG enzyme superfamily, free amino acid hydroxylases are attractive candidates for engineering new reactions.^23–26^ Because Fe(II)/αKG amino acid hydroxylases already have catalytic machinery to interact with amine and carboxylate functional groups in amino acids, we hypothesized that they might have promiscuous activity for substrates containing only an amine or only a carboxylate. These molecules are important feedstocks for early-stage oxyfunctionalization reactions in multi-step syntheses. Selectivity for these reactions has been historically difficult to achieve with traditional transition metal catalysis, and a biocatalytic process could offer improved regio- and stereoselectivity.^32,33^

We initially screened a panel of 12 Fe(II)/αKG amino acid hydroxylases for the ability to hydroxylate free carboxylates (Figure 1). We chose a set of candidate carboxylate substrates (**1-3**) that are structurally analogous to the native amino acid substrates L-pipecolic acid, L-proline, and L-leucine. We used whole-cell biocatalysis and liquid chromatography-mass spectrometry (LC-MS) to detect products. We confirmed that the native amino acid reaction products are detectable with all 12 members of the enzyme panel (Table S6). We then screened for promiscuous activity with carboxylates, and observed that one enzyme, tP4H,^34^ has detectable activity with substrate **1** (Figure 1). The total turnover number (TTN) with this substrate was ∼5 after 24 hr incubation with 10 μM enzyme, ∼130-fold lower than the TTN for the corresponding native amino acid substrate. The reaction of tP4H with substrate **1** gives the *trans* product with a *d.r.* of 4:1 (Figure S4). tP4H produces exclusively *trans* product with its native substrate L-pipecolic acid,^34^ suggesting that the free carboxylate substrate **1** and the native substrate are positioned similarly in the enzyme active site with respect to the iron center. To confirm that tP4H and not a contaminating enzyme was responsible for the observed non-native activity, we mutated active site residues that are involved in Fe(II), substrate, or αKG binding. Because tP4H does not have an experimental structure, we identified these active site residues using an Alphafold2^35^ model (Figure S9) and comparisons to structures of the highly homologous Fe(II)/αKG enzyme GriE.^36,37^ In all cases, active site mutations produced activity decreases for the non-native substrate **1** (Figure 2). We also observed increased product yield with increasing wild-type tP4H concentration (Figure 2). Together these results confirm that tP4H is responsible for the non-native reaction.

**Figure 1.**
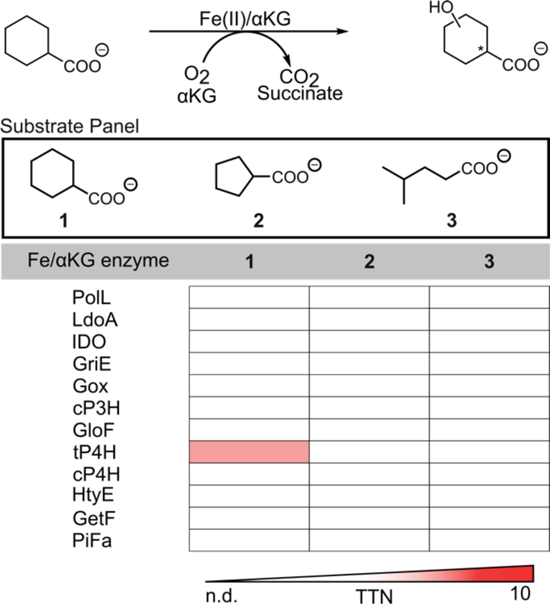
Initial whole-cell reaction screen data with a panel of Fe(II)/αKG amino acid hydroxylases and free acid substrate analogues. Reactions were performed in whole cell from 50 mL expression cultures where whole cell volume was 1/20^th^ the expression volume. Reactions were carried out in MOPS (pH 7.0, 50 mM) with 20 mM substrate, 60 mM αKG (as disodium salt), 1 mM ferrous ammonium sulfate, and 1 mM L-ascorbic acid.

**Figure 2.**
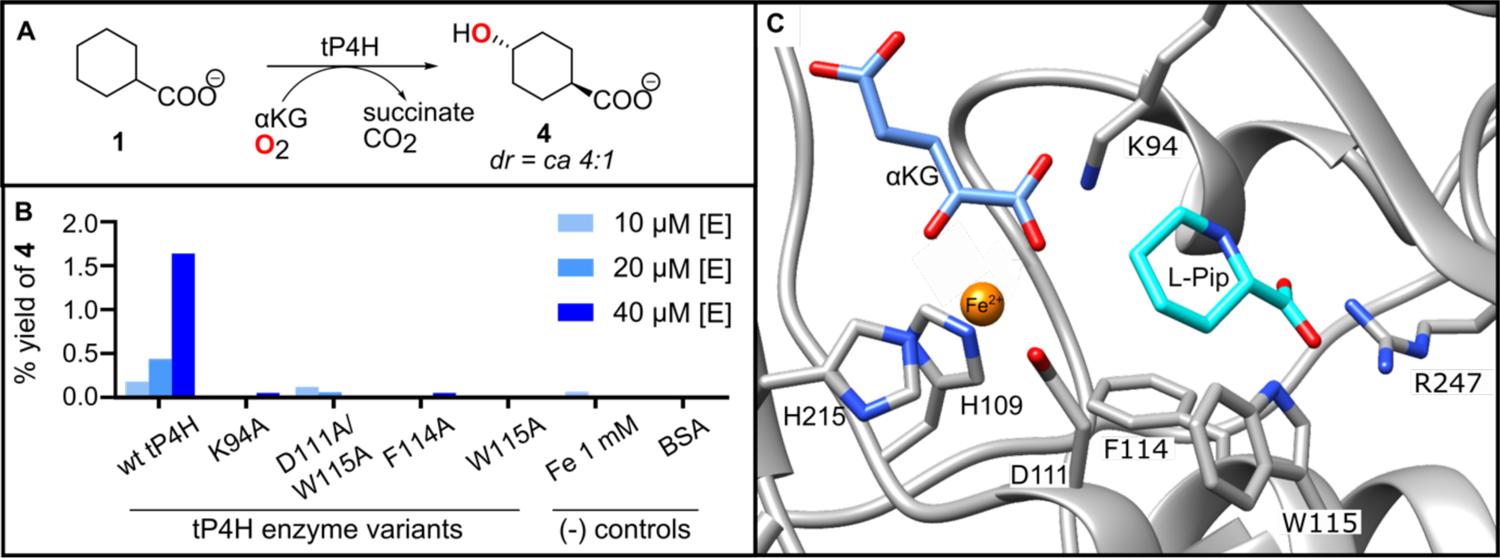
Validation of tP4H activity with free acid **1.** (A) Reaction of tP4H with substrate **1** to form *trans*-4-hydroxycyclohexane carboxylic acid **4**. (B) Yield of **4** after reaction of **1** with tP4H variants, as well as negative control reaction with Fe(II) and bovine serum albumin (BSA). The Fe 1 mM control was run in the absence of added enzyme. Purified enzyme concentration was varied between 10-40 μM with 20 mM **1**, 40 mM αKG, 1 mM ferrous ammonium sulfate, and 1 mM L-ascorbic acid in MES buffer (50 mM, pH 6.8). Reactions were carried out for 24 hours at 25 °C and quantified with analytical LC-MS. (C) Structural model of the tP4H showing key active site residues. Fe(II), αKG, and L-Pipecolic acid were modeled in Chimera.^38^

### Stabilization of tP4H with ProteinMPNN

To improve tP4H activity towards substrate **1**, we began a directed evolution campaign but quickly encountered limitations due to poor enzyme stability. First, we found that tP4H variants were difficult to express and purify due to enzyme insolubility. Additionally, we found that the parent wild-type tP4H enzyme loses activity with time (Figure 3). These observations are consistent with prior reports on tP4H behavior.^34^ It is possible for enzyme stability to improve during directed evolution, whether by selecting more stable variants during each round or by chance. For example, the evolved Fe(II)/αKG PsEFE had slight stability improvements compared to wild-type, despite the researchers not selecting for improved stability.^39^ In another case, the Fe(II)/αKG UbP4H was successfully screened for improved stability after initial screening rounds destabilized the enzyme.^27^ However, the benefits of starting with a highly stable enzyme for directed evolution are well-established.^12–14^ A stabilized protein scaffold could potentially increase the population of active, properly-folded protein or provide access to other mutants that otherwise do not fold.

**Figure 3.**
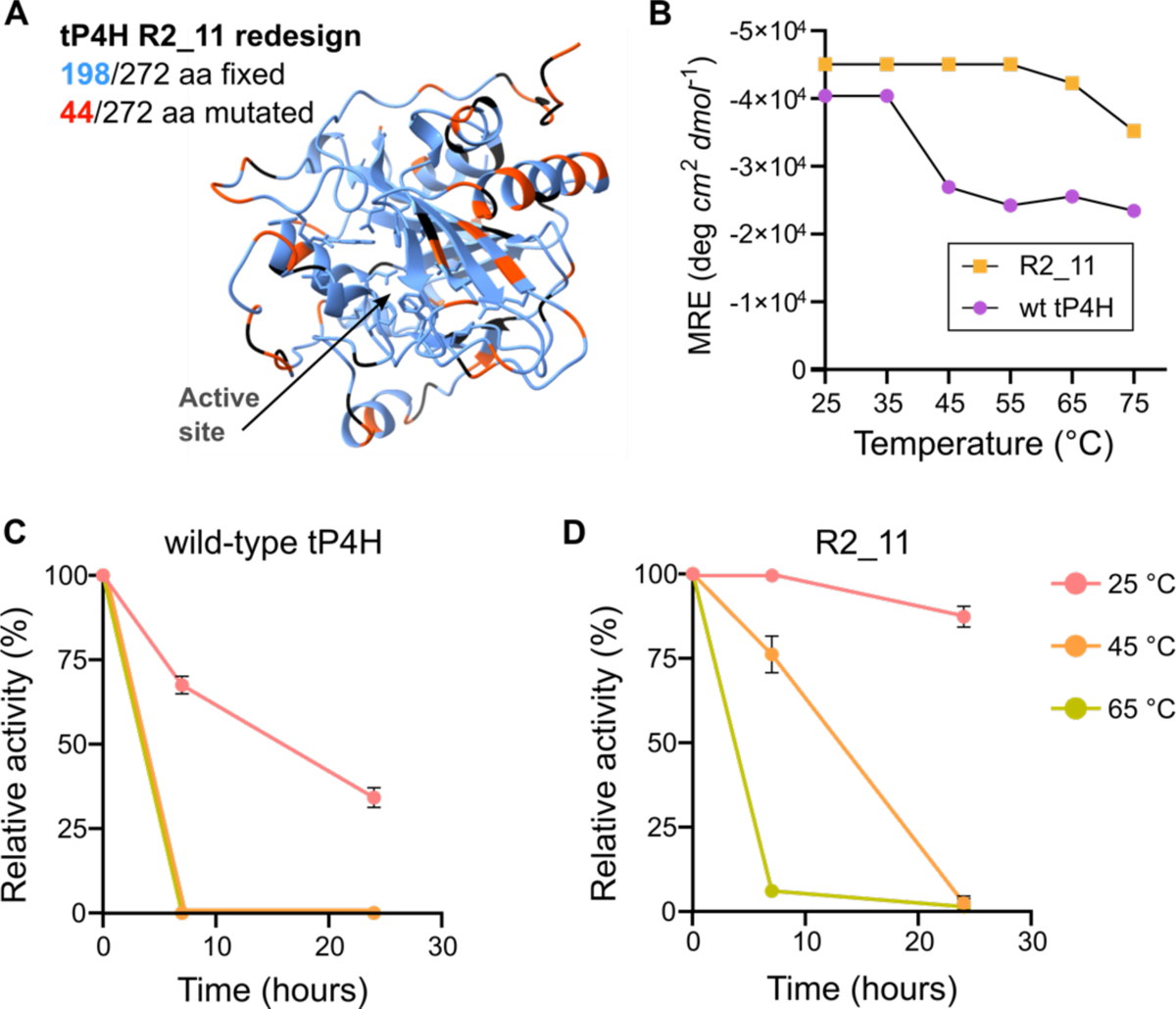
Stability and activity of wild-type tP4H and ProteinMPNN design R2_11. (A) tP4H structure (Alphafold2 model, Figure S9) color-coded to show sites fixed in the design process (blue, Supplementary spreadsheet – ProteinMPNN sequences_metrics) and sites mutated in the ProteinMPNN R2_11 redesign (orange-red). Sites colored black were neither fixed nor redesigned in the R2_11 variant. Side chains for first shell active site residues (Table S4) are shown in blue. (B) Temperature-dependent CD spectroscopy of wild-type tP4H and R2_11. (C) Activity-stability analysis of wild-type tP4H. (D) Activity-stability analysis of R2_11. Relative activity was determined using PBP assay described in Supplementary Information.

We used the deep learning-based tool ProteinMPNN^29,30^ to generate stabilized variants of tP4H through sequence design. We first used ProteinMPNN to redesign the entire tP4H sequence. Unsurprisingly, we found that the predicted sequences eliminated key active site residues, which is likely to disrupt enzymatic activity (Figure S10). This behavior is consistent with the well-established propensity for catalytic active site residues to be destabilizing.^3–10^ To preserve catalytic function, we fixed the active site residues in all subsequent design efforts. We defined the active site as any residues that contact the amino acid substrate, Fe(II), or the αKG cofactor, based on our Alphafold2 model (Figure S9) and comparisons to the structure of the closely-related enzyme GriE bound to L-leucine.^37^ Because other residues throughout the protein could also be important, we also tested four additional strategies using either sequence conservation or distance metrics. To identify important conserved residues, we constructed a multiple sequence alignment (MSA) and selected tP4H residues conserved in at least 35%, 70%, or 95% of sequences (Methods and Table S4). Alternatively, we fixed any residues with side chains within a 10 Å sphere from the substrate binding pocket. Using these five starting points (fix active site only, active site + 35%/70%/95% conservation, active site + 10 Å sphere), we generated 48 ProteinMPNN sequences per method and selected 4 each (20 total) for activity screens. Selection was based on calculated top-ranked Cα-RMSD values matched to the input tP4H structure. We obtained only one variant that had any detectable activity, with catalytic efficiency ∼35-fold lower than wild-type tP4H (Figure S11). This variant was designed from the sequence where >35% conserved residues are fixed, which constrains more residues than the >70% or >95% cutoffs. This result suggests that even weakly conserved residues may need to be fixed to maintain activity. Further analysis of the variant with detectable activity revealed a ∼3.5-fold increase in the *K*_M_ for the αKG cofactor (Figure S11). We identified two residues in proximity to αKG, L228 and V230, that were mutated in the redesigned sequence. These sequence changes may have contributed to improved stability at the expense of cofactor binding and positioning, leading to the decrease in activity. Notably, L228 and V230 were fixed in the designs generated from fixed active site + 10 Å sphere, but none of these designs had detectable activity. Taken together, these findings suggest that additional criteria will be needed to identify critical functional residues that should be fixed prior to sequence redesign.

To generate stabilized variants that maintain catalytic activity, we performed another set of ProteinMPNN sequence redesigns with three new strategies to fix important residues. In each case, we fixed the active site as defined above plus residues L228 and V230. For the first approach, we fixed all residues at tP4H positions conserved in 35% of the MSA. We chose this cutoff because it was the only one from our initial set that produced a stabilized variant with any detectable activity, and we expected that fixing L228 or V230 could further improve these designs. For the second and third approaches, we identified highly conserved positions regardless of whether the wild-type tP4H residue is the most highly conserved amino acid. These strategies were based on previous work suggesting that more stringent constraints are necessary to maintain activity in ProteinMPNN redesign.^30^ Every tP4H amino acid position was ranked based on the % conservation of the most frequent amino acid present in the MSA, and the top 50% or 70% were fixed as the wild-type tP4H residue. Together with fixed active site residues, these criteria resulted in 148/272 (54%) or 198/272 (73%) fixed residues across the entire 272 amino acid protein. Using these three strategies, we selected 32 designs each of 48 generated for a total of 96 sequences (Supplementary spreadsheet – ProteinMPNN sequences_metrics). Of these designs, 69 expressed detectable quantities of protein by SDS-PAGE and 11 had detectable activity above background for the native substrate. For the active enzymes we proceeded to measure thermostability and kinetic parameters for the native L-pipecolic acid substrate and the promiscuous carboxylate substrate **1**. The variant with the highest *k*_cat_ for L-pipecolic acid was R2_11 (Table S7), with a *k*_cat_ of 0.16 s^-1^ compared to 0.17 s^-1^ for wild-type. R2_11 was designed from the method where the top 70% ranked conserved residues were fixed, and had 44 designed mutations compared to the wild-type sequence (Figure 3). R2_11 has modestly slower (∼3-fold) non-native carboxylate hydroxylase activity compared to wild-type tP4H and exhibits a 20 °C increase in thermostability by temperature-dependent circular dichroism (CD) spectroscopy (Figure 3). When activity is measured as a function of time, R2_11 maintains activity over a timescale of days, which is a substantial improvement compared to wild-type tP4H (Figure 3).

### Stabilization of GriE with ProteinMPNN

After successfully identifying sequence constraints for ProteinMPNN-mediated stabilization of tP4H while maintaining catalytic function, we evaluated whether the same approach would be effective with a second Fe(II)/αKG enzyme, GriE. This enzyme could benefit from stabilization because, although it expresses well and is soluble, it loses activity at room temperature over 24 hours (Figure S12). As with tP4H, we fixed the top 70% ranked conserved residues along with catalytic residues identified from the GriE crystal structure (Supplementary spreadsheet – ProteinMPNN sequences_metrics).^37^ We generated 32 redesigned sequences and found that 29 were expressed as soluble enzyme, and 27 showed activity with the GriE native substrate L-leucine. The top design based on stability and kinetic parameters, GM_A9, showed a catalytic efficiency (*k*_cat_/*K*_M_) 4-fold lower than wild-type GriE (Figure S12 & S13). One design, GM_A11, had a 2-fold faster initial rate with L-leucine compared to GM_A9 but this design was unstable by temperature-dependent CD and thus was not chosen for further analysis. For GM_A9, the decrease in catalytic efficiency arises from a lower *k*_cat_ of 0.38 s^-1^ compared to our measured *k*_cat_ of 2.1 s^-1^ for wild-type GriE. A decrease in catalytic efficiency is not surprising given that increased stability could reduce conformational flexibility and negatively impact catalytic function.^4,6,7^

We next screened the stabilized GriE redesign GM_A9 for substrate promiscuity. Previously, wild-type GriE has been shown to accept substrates with increased substrate chain lengths but has weaker activity towards substrates with substitution at C3.^40^ We chose two previously identified non-native substrates to test: L-norleucine and L-allo-isoleucine. L-norleucine was chosen as a representative substrate with increased chain length compared to L-leucine. L-allo-isoleucine was chosen because it has a methyl group substitution at C3. We observed detectable activity with L-norleucine but not for L-allo-isoleucine (Table S8). Similar to wild-type GriE, the GM_A9 variant maintained a preference for the extended chain L-norleucine substrate over the C3-substituted L-allo-isoleucine substrate. The GM-A9 reactions with L-leucine and L-norleucine were 11- and 4-fold slower than wild-type GriE reactions, respectively (Table S8). These results suggest that our ProteinMPNN protocol can be readily applied to other Fe(II)/αKG enzymes to stabilize proteins while maintaining synthetically-relevant catalytic function that can be a foothold for further optimization by directed evolution.

### Directed evolution of wild-type tP4H for carboxylate C-H hydroxylation activity

We next sought to improve the non-native carboxylate hydroxylase activity through directed evolution. We prioritized tP4H because carboxylate hydroxylase activity was detectable in both the wild-type and ProteinMPNN-stabilized variant, which allows for direct comparisons. We conducted three rounds of directed evolution for both enzymes by varying first- and second-shell substrate binding residues identified in the active site from our Alphafold2 structural model. We defined the first shell as any residues that contact the amino acid substrate, based on comparisons to the structure of ligand-bound GriE.^37^ We defined the second shell as any residues that make contacts with first shell residues. We used the 22c-trick method for single site-saturation mutagenesis at each target position.^41^

We first performed directed evolution with wild-type tP4H. For the first screening round, we chose three tP4H active site residues based on their potential role in substrate specificity: H58, F114, and L174. Based on our tP4H structural model (Figure S9B), H58 likely contacts the amine of native amino acid substrates and is presumably not needed or detrimental for carboxylate substrates that lack an amine. F114 likely provides a substrate hydrophobic contact, and L174 is part of a loop that could affect substrate binding. We screened whole cell biocatalysis reactions in 96 well plates for improved TTN and 80% *trans* selectivity in reactions with substrate **1**. Based on production of hydroxylated product **4**, the top 5% of mutants were chosen for validation with purified enzymes. We obtained several variants with modest activity improvements, and the best performer was mutant H58L with a TTN of 7 (Figure 4A). In a second round starting from H58L, we rescreened mutants at F114 and L174 and screened an additional 14 first and second shell residues (Table S1). We identified the improved variant H58L/W170Q with a TTN of 15. In a third round starting from H58L/W170Q, we screened 9 residues that showed activity increases in previous rounds and identified the improved variant H58L/W170Q/E118K with a TTN of 31 (Figure 4A & Table S1). Overall, after three rounds of directed evolution for improved carboxylate hydroxylase activity with wild-type tP4H we obtained a 6-fold improvement in TTN and maintained >80% selectivity for the *trans* reaction product (Figure S4).

**Figure 4.**
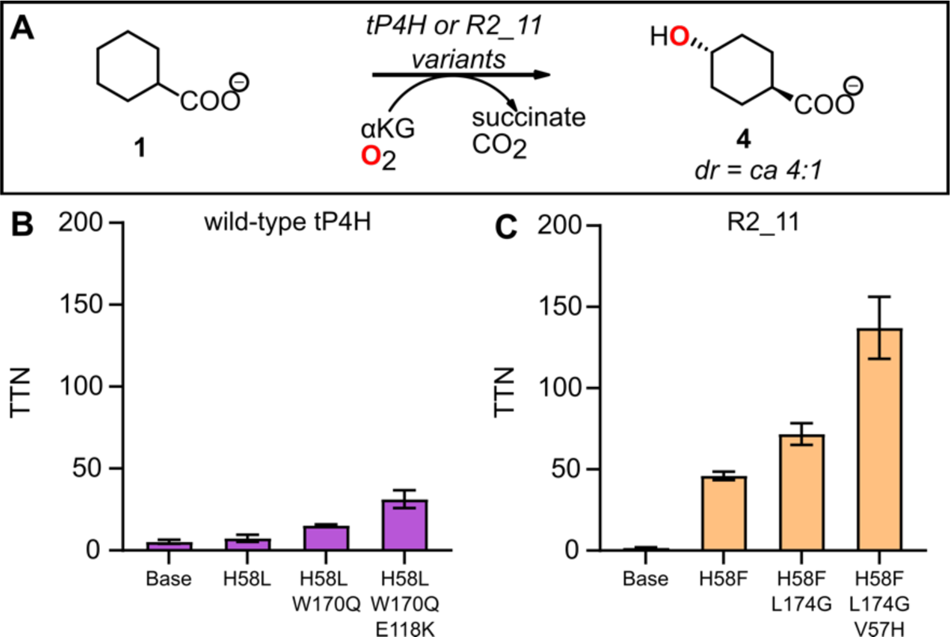
(A) Reaction scheme for C-H hydroxylation of substrate **1** with tP4H, R2_11, and associated variants. (B) Directed evolution of wild-type tP4H. (C) Directed evolution of stabilized variant R2_11. Reactions were carried out for 24 hr at 25 °C using purified enzyme (10-20 μM) in MES buffer (50 mM, pH 6.8), with 20 mM cyclohexane carboxylic acid **1**, 40 mM αKG, 1 mM ferrous ammonium sulfate, and 1 mM ascorbic acid. Concentration of **4** in quenched reaction samples was quantified by analytical LC-MS analysis.

### Directed evolution of ProteinMPNN redesign R2_11 for carboxylate C-H hydroxylation activity

To optimize carboxylate hydroxylase activity in ProteinMPNN-stabilized tP4H, we conducted directed evolution using a similar strategy to our approach with wild-type tP4H, with minor modifications. In the first round of site saturation mutagenesis, we started with a larger pool of 19 first- and second-shell residues including the three sites from prior round one (H58, F114, and L174) and the 16 additional sites from prior round 2 (Table S1). As before, we screened whole cell biocatalysis reactions for improved TTN and >80% *trans* selectivity with carboxylic acid substrate **1**. The top hit was H58F, which is the same position but a different mutant than the previous round one winner, H58L. R2_11_H58F displayed a 27-fold increase in TTN relative to the parent R2_11 (Figure 4B). This effect is substantially bigger than the <2-fold improvement obtained with H58L relative to wild-type tP4H. Notably, the 27-fold increase in a single round from stabilized R2_11 was already larger than the total 6-fold improvement from three rounds of directed evolution from wild-type tP4H. Given the strong performance of the H58F mutant, we also evaluated its effect in the wild-type tP4H background and observed a small, <2-fold increase in TTN, similar to the effect of H58L on wild-type tP4H (Figure S14). Thus, the strong, 27-fold improvement with the H58F mutant depends on the context of the stabilized R2_11 backbone.

In a second round of screening from R2_11_H58F, we selected the 18 residues that were screened in prior round two from wild-type tP4H (Table S1). We retained this large pool of residues to ensure a direct comparison to the tP4H directed evolution workflow. This round identified improved variant L174G. R2_11_H58F/L174G has a TTN of 72. This TTN is a 1.6-fold improvement from parent R2_11_H58F and outstrips any variant obtained from the wild-type tP4H backbone (Figure 4B). In a third round of screening from R2_11_H58F/L174G, we selected 9 residues that were screened in prior round 3 from wild-type tP4H (Table S1). This round identified the improved variant V57H with a TTN of 138, a 1.7-fold improvement from the previous round.

Overall, the ProteinMPNN-stabilized tP4H directed evolution campaign produced an 80-fold improvement in TTN from the base R2_11 redesign, compared to a modest 6-fold improvement in the wild-type tP4H evolutionary trajectory. Although the R2_11 parent starts ∼3-fold slower than wild-type tP4H, the much larger improvement over three rounds of directed evolution produced an R2_11 triple mutant with a 4.5-fold higher TTN than the triple mutant obtained from wild-type tP4H (Figure 4).

In addition to a more efficient directed evolution trajectory, the R2_11 triple mutant maintains high stability relative to the wild-type tP4H and associated variants from directed evolution (Figure 5A, Figure S15). Higher stability allows reactions to be run more efficiently, both at higher temperatures and for less time. For example, after 6 hours at 35 °C, the R2_11 triple mutant reaches a mean TTN of 142 for the non-native reaction with carboxylate **1** to form product **4** with 4:1 selectivity for the *trans* reaction product (Figure 5B). The TTN after 6 hours at 35 °C is comparable to the TTN after 24 hours at 25 °C. In contrast, the tP4H triple mutant shows a slight decrease in TTN at 35 °C, likely due to enzyme instability at higher temperatures (Figure 5B). The stability profile of the R2_11 triple mutant suggests that this enzyme will be more robust towards further engineering compared to the tP4H triple mutant. Future engineering efforts with the R2_11 mutant could include improvements to key reaction metrics like turnover, selectivity, and increased substrate scope.

**Figure 5.**
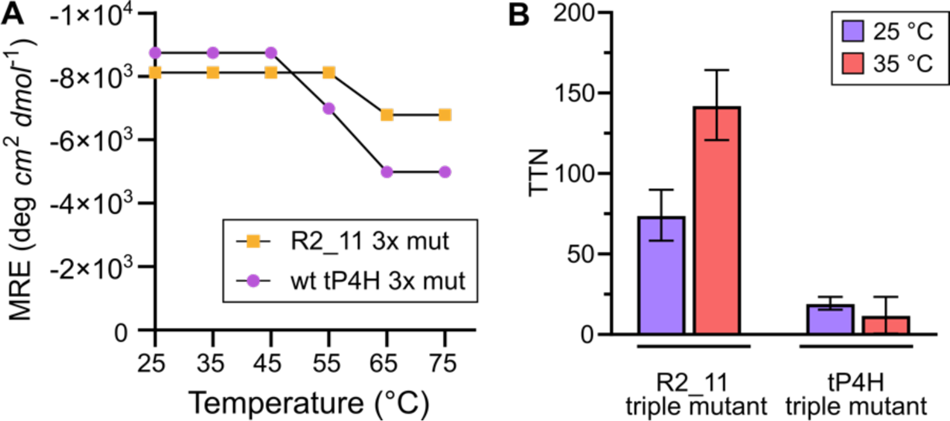
A) Temperature dependent CD of the R2_11 and tP4H triple mutants. B) TTN for formation of **4** (*d.r.* 4:1, Figure S4) with the R2_11 and tP4H triple mutants at two different temperatures. Reactions were carried out for 6 hrs. at 25 °C and 35 °C using purified enzyme (15 μM) in MES buffer (50 mM, pH 6.8), with 20 mM cyclohexane carboxylic acid **1**, 40 mM αKG, 1 mM ferrous ammonium sulfate, and 1 mM ascorbic acid.

## Conclusions

Directed evolution is a powerful tool to engineer enzymes for new-to-nature reactions. However, many enzyme starting points for evolution may lack the stability required to reach user-defined optimum fitness after multiple rounds of mutagenesis. Here we show that the deep learning-based tool ProteinMPNN can be used to stabilize the Fe(II)/αKG enzyme superfamily members tP4H and GriE with straightforward sequence constraints to maintain catalytic activity. Consistent with previous results using ProteinMPNN,^30^ the top tP4H design was identified by using the most conservative of our chosen methods for fixing residues during sequence redesign. Applying the same method to the related enzyme GriE readily produced stabilized variants with catalytic activity.

Wild-type and redesigned tP4H both exhibit novel reactivity towards remote C-H hydroxylation of a free carboxylic acid substrate. We directly compared evolutionary trajectories of wild-type tP4H with the stabilized variant R2_11 and demonstrated superior performance of the stabilized redesign variant. Future work will determine if this design method is generalizable to optimize directed evolution for other enzymes and enzyme families. User-friendly deep learning-based tools like ProteinMPNN and MutCompute are rapidly emerging,^22,29,30^ and our work suggests that these tools should be routinely incorporated into enzyme engineering workflows to efficiently optimize catalytic fitness for new biocatalysts.^31^

## Supplementary Information

Materials, experimental and analytical methods, compound characterization data (Figure S4-S7), and enzyme characterization data (Figure S3, Figure S8, Figure S11-S15 and Tables S6-S8) (PDF). Accession numbers for all Fe(II)/αKG enzymes used in this work, full nucleotide and amino acid sequences for all reported wild-type and enzyme variants, oligonucleotide sequences for cloning and for all ProteinMPNN designs, ProteinMPNN design screen criteria results (XLSX).

## Acknowledgements

We thank Dr. Wolfgang Hüttel at the University of Freiburg and Hans Renata at Rice University for the donation of the wild-type tP4H and GriE expression vectors, respectively, and for their advice on tP4H and GriE reactions in various formats. We also thank Jonathan Zhang and Susanna Vazquez Torres for their early contributions to Fe(II)/αKG reaction screening, and Dr. Martin Sadilek in University of Washington Mass Spectrometry Facility for his continued support and helpful advice in analytical method development. This work was supported by U.S. National Institutes of Health grants T32 GM008268 (B.R.K., J.L.C.) and R35 GM124773 (J.G.Z.), and by the Open Philanthropy Project Improving Protein Design Fund (K.H.S., D.B.).

