## Supplementary Information for "Computational stabilization of a non-heme iron enzyme enables efficient evolution of new function"

Table of Contents

MATERIALS AND METHODS ... 4

Starting materials and product markers ... 4

Analytical instrumentation ... 4

General procedure for FMOC-Cl derivatization of Fe/αKG amino acid substrates and products ... 4

DNA and protein sequences for wild-type and base MPNN designs ... 4

Materials ... 4

Plasmids and Proteins ... 5

Protein overexpression ... 8

Protein purification ... 8

Site-saturation mutagenesis and library screening ... 10

Cloning ... 10

Library Expression ... 11

Library reactions in whole-cell ... 12

Library analysis ... 13

Library hit selection ... 13

Library hit validation ... 14

Scale-up biocatalytic reaction with R2_11 H58F/L174G/V57H and product isolation ... 14

Continuous fluorescence-based assay for Fe(II)/αKGs ("PBP assay") for initial rates measurements and Michaelis-Menten analyses ... 15

General set-up ... 15

Reagents and sources ... 18

Purification of succinyl-CoA synthetase ... 18

Assay validation ... 19

ProteinMPNN computational sequence redesign ... 20

ProteinMPNN multiple sequence alignment (MSA) input ... 20

Fixed Residue Selection ... 20

ProteinMPNN Design of tP4H and GriE ... 21

Design cloning and screening ... 22

EXPERIMENTAL DATA ... 27

SDS-PAGE of tP4H variants and ProteinMPNN redesigns ... 27

LC-MS characterization of substrate 1 hydroxylation products and calibration curves ... 28

NMR spectra of isolated product 4 and product markers ... 30

LCMS traces for reactions of GriE and GM_A9 ... 31

### MATERIALS AND METHODS

#### Starting materials and product markers

All starting materials and product markers were used as received from commercial suppliers, unless otherwise noted.

#### Analytical instrumentation

HPLC-UV-MS analysis for native enzyme reactions after Fmoc-Cl derivatization was performed with an Agilent 1100 LC equipped with a Zorbax C18 column (4.6 x 100 mm, 2.1 µm ID) with DAD detection connected to a Bruker IonTrap MS. HPLC-MS analysis for site-saturation mutagenesis library screening was performed on a Waters Xevo-TQ equipped with a Waters BEH-Amide column (2.1 x 50mm 1.7 µm, plus guard column). Continuous fluorescence measurements were collected on a PerkinElmer plate-reader. Circular dichroism measurements were collected with a JASCO J-1500 instrument using a 1 mm pathlength cuvette.

#### General procedure for FMOC-Cl derivatization of Fe/αKG amino acid substrates and products

To 100 µL of quenched biocatalytic reactions was added 100 µL of 200 mM sodium borate (pH 8.0) and 20 µL of 10 mM 9-fluorenylmethyl chloroformate (Fmoc-Cl) in acetone. After vortexing for 60 seconds, 20 µL of 100 mM 1-adamantanamine solution in acetone was added to stop the reaction followed by vortexing for 60 seconds. 50 µL of the resultant derivatized mixture was added to a 50:50 mixture of water and acetonitrile and submitted for LC/MS analysis. LC/MS analysis for derivatized amino acid products was carried out using an Agilent 1100 LC connected to a DAD detector and a Bruker IonTrap mass spec in positive mode. Separation was carried out on a Zorbax C18 column (4.6 x 100 mm, 2.1 µm ID) with an acid modified ACN/water gradient.

#### DNA and protein sequences for wild-type and base MPNN designs

##### Materials

DNA oligonucleotides were purchased from IDT. PCR was carried out using the Phusion® High-Fidelity PCR kit (New England Biolabs). Cloning of DNA fragments into linearized vectors was carried out using In-Fusion Snap Assembly Master Mix (Takara). All DNA and amino acid sequences used in this work are supplied in a supplementary Excel document.

### Plasmids and Proteins

#### *tP4H sequences*

The gene encoding *Dactylosporangium sp.* L-proline-4-hydroxylase *tP4H* (Uniprot ID O06499) was contained within a pET-28a(+) vector with an N-term 6xHis-tag. This plasmid, hereby referred to as pET28a-tP4H, was generously donated by Prof. Wolfgang Hüttel's lab at the University of Freiburg, Germany.[1] ProteinMPNN designs were purchased as IDT eBlocks that were cloned directly into a pET-28a(+) vector backbone between NcoI and XhoI for initial screening. All other DNA sequences are supplied in a supplementary Excel spreadsheet. For MPNN design directed evolution, the gene encoding R2_11 was cloned into a pET-22b(-) vector with carbenicillin resistance between XbaI and XhoI.

*tP4H* DNA sequence and In-Fusion overhangs for pET-28a(+).15 bp In-Fusion overhangs are bolded, and restriction sites are highlighted in red:

**5′aggagatataccatg**ggcagcagccatcatcatcatcatcacagcagcggcctggtgccgcg
cggcagccatatggctagcatgactggtggacagcaaatgggtcgggatccgaattcgagctcc
gtcgacggtatcgataagcttgatatcgaattcctgcagcccgacATGCTGACCCCGACCGAAC
TGAAACAGTACCGTGAAGCTGGTTACCTGCTGATCGAAGACGGTCTGGGTCCGCGTGAAGTTGA
CTGCCTGCGTCGTGCTGCTGCTGCTCTGTACGCTCAGGACTCTCCGGACCGTACCCTGGAAAAA
GACGGTCGTACCGTTCGTGCTGTTCACGGTTGCCACCGTCGTGACCCGGTTTGCCGTGACCTGG
TTCGTCACCCGCGTCTGCTGGGTCCGGCTATGCAGATCCTGTCTGGTGACGTTTACGTTCACCA
GTTCAAAATCAACGCTAAAGCTCCGATGACCGGTGACGTTTGGCCGTGGCACCAGGACTACATC
TTCTGGGCTCGTGAAGACGGTATGGACCGTCCGCACGTTGTTAACGTTGCTGTTCTGCTGGACG
AAGCTACCCACCTGAACGGTCCGCTGCTGTTCGTTCCGGGTACCCACGAACTGGGTCTGATCGA
CGTTGAACGTCGTGCTCCGGCTGGTGACGGTGACGCTCAGTGGCTGCCGCAGCTGTCTGCTGAC
CTGGACTACGCTATCGACGCTGACCTGCTGGCTCGTCTGACCGCTGGTCGTGGTATCGAATCTG
CTACCGGTCCGGCTGGTTCTATCCTGCTGTTCGACTCTCGTATCGTTCACGGTTCTGGTACCAA
CATGTCTCCGCACCCGCGTGGTGTTGTTCTGGTTACCTACAACCGTACCGACAACGCTCTGCCG
GCTCAGGCTGCTCCGCGTCCGGAATTCCTGGCTGCTCGTGACGCTACCCCGCTGGTTCCGCTGC
CGGCTGGTTTCGCTCTGGCTCAGCCGGTTTAAgatgggggatccactagttctagagcggccgc
ac**ctcgagcaccaccacc 3′**

tP4H protein sequence:

MLTPTELKQYREAGYLLIEDGLGPREVDCLRRAAAALYAQDSPDRTLEKDGRTVRAVHGCHRRD
PVCRDLVRHPRLLGPAMQILSGDVYVHQFKINAKAPMTGDVWPWHQDYIFWAREDGMDRPHVVN
VAVLLDEATHLNGPLLFVPGTHELGLIDVERRAPAGDGDAQWLPQLSADLDYAIDADLLARLTA
GRGIESATGPAGSILLFDSRIVHGSGTNMSPHPRGVVLVTYNRTDNALPAQAAPRPEFLAARDA
TPLVPLPAGFALAQPV*

272 amino acids

<u>*R2_11 base* DNA sequence and In-Fusion overhangs for pET-28a(+). 15 bp In-Fusion overhangs are bolded, and restriction sites are highlighted in red:</u>

**5′aggagatataccatg**ggcagcagccatcatcatcatcatcacagcagcggcctggttggcag
cATGCTGACCGATGAAGAACTGAAACGCTATAACGAACTGGGCTATCTGCTGATTGAAGATGGC
CTGGGCCCGGAAGAAGTGGCAGTTTTACGTCGCGCGGCGGATGAACTGTTTGCGGAAGATAGCC
CGGATCGCACCCTGGAGAAAGATGGCGTTACCGTGCGCAGCGTGCATGGTTGCCATCGTCGTAA
CCCAGTGTGCGCAGATTTAGTTCGTCATCCGCGTCTGCTGGGTCCGGCGCAGCAGATTTTAAGC
GGCGAAGTGTATGTGCATCAGTTTAAAATTAACGCCAAAGCGCCGATGCGTGGCGATGTGTGGC
CGTGGCATCAGGATTATATCTTTTGGAACCGCGAAGATGGCATGGATAAACCGCATGTGGTGAA
CGTGGCGGTGCTGCTGGATGAAGCGACCCATCTGAACGGCCCGTTACTGTTTGTGCCGGGCACC
CATGAACTGGGTGAAATTGATGTTGCGCGCCGTAATCCACCGGGCGATGGTCCGGATCAATGGT
TACCGCAGCTGAGCGCGGATCTGGATTATGCGATTGATGATGATCTGCTGGCGACCCTGACCGA
TGGTCGCGGCATTGATTCTGCAACCGGCAAAGCGGGCAGCATTCTGCTGTTTGATAGCCGCATT
GTGCATGGCAGTGGTCGCAACATGAGCCCATTTCCGCGTCGTGTGGCGCTGGTGACCTATAACC
GCACCGATAACGCCTTACCGGAACAGGATAATCCGCGCCCGGAATTTCTGGCAGCGCGTGATGC
AACCCCATTAACCCCGCTGCCGGAAGGCTTTCGTCTGGCGGATCCACCGTAAc**tcgagcaccac**
**cacc 3′**

<u>*R2_11 base* protein sequence:</u>

MLTDEELKRYNELGYLLIEDGLGPEEVAVLRRAADELFAEDSPDRTLEKDGVTVRSVHGCHRRN
PVCADLVRHPRLLGPAQQILSGEVYVHQFKINAKAPMRGDVWPWHQDYIFWNREDGMDKPHVVN
VAVLLDEATHLNGPLLFVPGTHELGEIDVARRNPPGDGPDQWLPQLSADLDYAIDDDLLATLTD
GRGIDSATGKAGSILLFDSRIVHGSGRNMSPFPRRVALVTYNRTDNALPEQDNPRPEFLAARDA
TPLTPLPEGFRLADPP*

272 amino acids

***GriE sequences***

The gene encoding *Streptomyces sp.* L-leucine hydroxylase *GriE* (Uniprot ID A0A0E3URV8) was contained within a pET-22b(-) vector with a C-term 6xHis-tag. This plasmid, hereby referred to as pET22b-GriE, was generously provided by Prof. Hans Renata (Rice University, USA).[2] ProteinMPNN designs were purchased as IDT eBlocks that were cloned directly into a pET-22b(+) vector backbone between XbaI and XhoI. All other DNA sequences are supplied in a supplementary Excel spreadsheet.

*GriE* DNA sequence and In-Fusion overhangs for pET-22b(-).15 bp In-Fusion overhangs are bolded, and restriction sites are highlighted in red:

**5′**
**acaattcccctctag**aaataatttgtttaactttaagaaggagatatacatATGCAGCTGACC
GCCGATCAGGTGGAAAAATATAAATCCGATGGATACGTTCTGCTTGAAGGCGCTTTCAGCCCAG
AAGAGGTTCACGTCATGCGTCAGGCACTGAAAAAGGATCAAGAAGTCCAAGGACCTCATCGGAT
TTTAGAAGAAGATGGTCGCACGGTCCGCGCACTGTATGCTAGTCACACAAGACAAAGCGTATTC
GATCAGTTGTCTCGCTCTGACAGATTACTGGACCCGCCACACAGCTGTTAGAGTGCGACTTAT
ACATTCACCAATTCAAGATTAATACTAAGCGCGCTTTTGGTGGAGATAGCTGGGCATGGCACCA
GGACTTTATCGTGTGGCGCGATACTGACGGTTTACCTGCCCCGCGTGCCGTCAATGTCGGCGTC
TTTTTATCGGACGTGACAGAGTTTAATGGGCCAGTGGTCTTTTTATCTGGCTCCCACCAGCGTG
GTACAGTGGAGAGAAAGGCACGGGAGACGTCACGCTCAGACCAGCATGTGGACCCGGATGATTA
TTCTATGACGCCAGCTGAGCTGAGCCAAATGGTGGAAAAACATCCAATGGTCTCCCCAAAGGCC
GCAAGCGGTTCTGTAATGTTGTTCCACCCAGAGATAATACACGGATCGGCACCAAACATCTCGC
CGTTTGCTCGTGACCTGTTGATCATTACATACAACGACGTGGCGAACGCCCCGAAACCGGCAGG
AGAACCACGCCCGGAATACGTTATCGGTCGTGATACTACGCCACTGGTGTCTCGCTCGGACCA
TTACACGAAGCAGCGGAGAGCCGGCTTGCCc**tcgagcaccaccacc 3′**

GriE protein sequence:

MQLTADQVEKYKSDGYVLLEGAFSPEEVHVMRQALKKDQEVQGPHRILEEDGRTVRALYASHTR
QSVFDQLSRSDRLLGPATQLLECDLYIHQFKINTKRAFGGDSWAWHQDFIVWRDTDGLPAPRAV
NVGVFLSDVTEFNGPVVFLSGSHQRGTVERKARETSRSDQHVDPDDYSMTPAELSQMVEKHPMV
SPKAASGSVMLFHPEIIHGSAPNISPFARDLLIITYNDVANAPKPAGEPRPEYVIGRDTTPLVS
RSGPLHEAAESRLA

270 amino acids

*GM_A9* sequence and In-Fusion overhangs for pET-22b(-).15 bp In-Fusion overhangs are bolded, and restriction sites are highlighted in red:

**5′**
**acaattcccctctag**aaataatttgtttaactttaagaaggagatatacatATGCAGCTGACC
GATGCGCAGGTGGAACAGTATAAAAGCGATGGCTATGTGCTGCTGGAAGGCGCGTTTAGCCCGG
AAGAAGTGGATATTATGCGTAAAGCGCTGGCGAAAGATGCGGAAGTTGAAGGCCCGCATCGCAT
TATGGAAGAAGATGGCAAAGCGGTGCGCGCGCTGTATGCGAGCCATAAACGCCAGAGCGTGTTT
GATCAGCTGAGCCGCAGCGATCGCCTGCTGGGCCCGGCGACCCAGCTGCTGGAATGCGATCTGT
ATATTCATCAGTTTAAAATTAACACCAAACGCGCGTTTGGCGGCAGCGCGTGGGCGTGGCATCA
GGATTTTATTGTGTGGCGCGATACCGATGGCCTGCCGGCGCCGCGCGCGGTGAACGTTGGCGTG
TTTCTGAGCGATGTGACCGAATTTAACGGCCCGGTGGTGTTCCTGAGCGGTAGCCATCAGAAAG
GCACCCTGGAACGCAAACGTCGCGCGACCAGCGTGAGCGATGAACATGTGGATCCGCGCGATTA
TAGCATGACCCCGGCGGAACTGGAGAAAATGGTGAAAGAACATCCGATGGTGAGCCCGAAAGCC
GCGAGCGGCAGCGTGCTGCTGTTTCATCCGGAAGTGATTCATGGCAGCTTTCCGAACATTAGCC
CGTTTGCGCGTGATCTGCTGATTATTACCTATAATGATGTGAACAACGCGCCGAAACCGGCGGG
CACCCCGCGCCCGGAATATGTGATTGGCCGTGATACCACCCCGCTGGTGAGCGAAAGCGGCCCG
CTGCATc**tcgagcaccaccacc 3′**

GM_A9 protein sequence:

MQLTDAQVEQYKSDGYVLLEGAFSPEEVDIMRKALAKDAEVEGPHRIMEEDGKAVRALYASHKR
QSVFDQLSRSDRLLGPATQLLECDLYIHQFKINTKRAFGGSAWAWHQDFIVWRDTDGLPAPRAV

NVGVFLSDVTEFNGPVVFLSGSHQKGTLERKRRATSVSDEHVDPRDYSMTPAELEKMVKEHPMV
SPKAASGSVLLFHPEVIHGSFPNISPFARDLLIITYNDVNNAPKPAGTPRPEYVIGRDTTPLVS
ESGPLH

262 amino acids – Note the C-term sequence “EAAESRLA” was excluded from ProteinMPNN design as it was not resolved in the crystal structure for GriE (PDB ID: 5NCI).[3]

### Protein overexpression

Chemically competent *E. coli* BL21(DE3) cells were transformed with plasmids containing *tP4H, GriE*, associated variants, and ProteinMPNN designs using a standard heat-shock protocol. Starter cultures of LB with the appropriate antibiotic were inoculated from a single *E. coli* colony on an agar plate encoding the protein of interest and grown overnight to stationary phase at 37 °C. Expression cultures of Terrific Broth supplemented with the appropriate antibiotic were inoculated with the starter cultures (2% v/v) and shaken at 37 °C at 200 rpm in a ThermoFisher Scientific MaxQ800 shaker. When expression cultures reached $OD_{600}$ of ~0.4-0.5 (typically 2-3 hours), they were cooled to 18 °C and protein expression was induced by addition of isopropyl β-D-1-thiogalactopyranoside (IPTG, 0.5 mM). Cultures were incubated at 18 °C and 200 rpm overnight (16-24 hours). Cells were pelleted by centrifugation (4 °C, 4000 rpm, 10 minutes). Cell pellets were resuspended in MES buffer (50 mM, pH 7.0) for whole-cell reactions. For reactions with lysate, whole-cell suspensions in MES buffer (50 mM, pH 7.0) were lysed by sonication on ice (70% power, 30 seconds, 1 second on/1 second off, repeated 3 times). Clarified lysate was prepared by centrifugation of lysed cells (4 °C 10,000 rpm, 15 min) and used directly in biocatalytic reactions.

### Protein purification

Large-scale purification: Harvested cell pellets from 1 L overexpression were resuspended on ice in 20 mL of Lysis Buffer (25 mM Tris pH 8.0, 150 mM NaCl, 5 mM Imidazole, 5% Glycerol) containing protease inhibitor (Pierce Protease Inhibitor Mini Tablets EDTA-free, ThermoFisher A32955). Resuspended cells were lysed by sonication (70% power, 30 seconds, 1 second on/1 second off, repeated 3 times), and clarified lysate was prepared by centrifugation (16,000 rpm, 20 min, 4 °C). Clarified lysate was incubated with Ni-NTA resin (3 mL bed volume in Econo-Pac® Chromatography Columns, Bio-Rad) for 30 min at 4 °C on a rocker. The resin was washed successively with 3 column volumes of each of the following buffers: Lysis buffer (25 mM Tris pH 8.0, 150 mM NaCl, 5 mM Imidazole, 5% Glycerol) NaCl wash buffer (25 mM Tris, pH 8.0, 1M NaCl, 5 mM imidazole,

5% glycerol), and Ni-wash buffer (25 mM Tris, pH 8.0, 150 mM NaCl, 15 mM imidazole, 5% glycerol). Enriched His-tagged protein was eluted from the resin by incubation for 10 min with Ni-elution buffer (5-10 mL, 25 mM Tris, pH 8.0, 150 mM NaCl, 250 mM, 10% glycerol). The eluted proteins were dialyzed into buffer suitable for enzymatic reactions (MES or MOPS 50 mM, pH 7.0, 10% glycerol). Protein concentration was determined by Bradford assay. Dialyzed protein used directly or was aliquoted into PCR or Eppendorf tubes, flash frozen in liquid nitrogen, and stored at -80 °C for further use. Figure S3 shows representative SDS gel samples for wild-type tP4H and R2_11 and associated variants from directed evolution rounds.

Small-scale purification: Harvested cell pellets from 25-50 mL expressions were resuspended on ice in 1-2 mL of lysis buffer (25 mM Tris pH 8.0, 150 mM NaCl, 5 mM Imidazole, 5% Glycerol) containing protease inhibitor (Pierce Protease Inhibitor Mini Tablets EDTA-free, ThermoFisher A32955). Resuspended pellets were lysed by sonication on ice in 1.5 mL Eppendorf tubes (70% power, 15 seconds with 1 second on/1 second off pulse, repeated 2 times). Clarified lysate was prepared by centrifugation (12,000 x g, 15 min, 4 °C). Clarified lysate was incubated with Ni-NTA resin in a fresh Eppendorf tube (300 µL of a 1:1 suspension of Ni-NTA resin to lysis buffer) at 4 °C for 30 min. Ni-NTA resin and His-tagged protein was spun down for 30 seconds at 12,000 x g, and supernatant was removed via pipetting. The resin bed was then washed in a similar manner with 3x1 mL washes of the following buffers: Lysis buffer (25 mM Tris pH 8.0, 150 mM NaCl, 5 mM Imidazole, 5% Glycerol) NaCl wash buffer (25 mM Tris, pH 8.0, 1M NaCl, 5 mM imidazole, 5% glycerol), and Ni-wash buffer (25 mM Tris, pH 8.0, 150 mM NaCl, 15 mM imidazole, 5% glycerol). Enriched His-tagged protein was eluted from the resin by 10-minute incubation with Ni-elution buffer (0.5-1 mL, 25 mM Tris, pH 8.0, 150 mM NaCl, 250 mM, 10% glycerol). Supernatant enriched with purified enzyme was removed after centrifugation to pellet Ni-NTA resin. Purified proteins were buffer exchanged into storage and reaction buffer (MES or MOPS 50 mM, pH 7, 10% glycerol) using an appropriately sized Zeba Spin Desalting Column with 7K MWCO. Protein concentration was determined by Bradford assay. Dialyzed protein was used directly or aliquoted into PCR or Eppendorf tubes, flash frozen in liquid nitrogen, and stored at -80 °C for further use.

### Site-saturation mutagenesis and library screening

#### Cloning

Site-saturation mutagenesis (SSM) was carried out using the 22c-trick method.[4] All oligos used can be found in a supplementary Excel spreadsheet. 22c-trick oligos were used in conjunction with carbenicillin forward and reverse oligos to amplify two plasmid fragments for In-Fusion assembly. The carbenicillin overlap was used as a positive control for correct plasmid assembly under antibiotic selection. All complete PCR reactions were treated with DpnI to eliminate template plasmid DNA. PCR products were isolated by gel electrophoresis using 1% agarose gels with visualization using Invitrogen SYBR Safe dye. Gel fragments were isolated after gel extraction (ThermoFisher GeneJet Gel Extraction kit). Each SSM library was constructed with two purified linear DNA fragments using In-Fusion assembly (Takara). After In-Fusion assembly, constructed plasmids were transformed into chemically competent NEB® Turbo Competent *E. coli* cells. Transformants were grown in 5 mL of LB-carbenicillin at 37 °C for 10-12 hrs to perform a quick quality control (QQC) check. After incubation, plasmid libraries were isolated after mini-prep (ThermoFisher GeneJet Plasmid Mini-prep kit) and sent for Sanger sequencing (Azenta). Confirmed libraries were then transformed into electrocompetent E. cloni EXPRESS BL21(DE3) cells (Lucigen, catalog #: 60300-2) and plated on LB-carb agar plates after a 1 hr outgrowth at 37 °C. The plated cells were incubated at 37 °C for 14-16 hr and stored at 4 °C until colony picking.

Sites chosen in each round of directed evolution for both wild-type tP4H and the R2_11 redesign are shown in Table S1.

Table S1

| tP4H Directed Evolution | | | MPNN R2_11 Directed Evolution | | |
|---|---|---|---|---|---|
| **Round 1** | **Round 2** | **Round 3** | **Round 1** | **Round 2** | **Round 3** |
| **H58** | V57 | V57 | V57 | V57 | **V57** |
| F114 | F93 | I95 | **H58** | F93 | I95 |
| L174 | I95 | P107 | F93 | I95 | P107 |
| | W106 | I113 | I95 | W106 | I113 |
| | P107 | **E118** | W106 | P107 | E118 |
| | Y112 | L174 | P107 | Y112 | L174 |
| | I113 | V231 | Y112 | I113 | V231 |
| | F114 | F250 | I113 | F114 | F250 |
| | W115 | | F114 | W115 | |
| | E118 | | W115 | E118 | |
| | D119 | | E118 | D119 | |
| | **W170** | | D119 | W170 | |
| | L174 | | W170 | **L174** | |
| | T232 | | L174 | T232 | |
| | V231 | | V231 | V231 | |
| | F250 | | T232 | F250 | |
| | | | F250 | | |

### Library Expression

Single colonies from the LB-carb agar plates for each SSM library were picked with sterile toothpicks and used to inoculate starter cultures using 0.5 mL LB-carb into 96-well deep well plates (Costar 3961). 70 colonies were picked for every library to ensure >95% library coverage. For a positive control, 5 colonies harboring plasmid encoding for the parent of the round were picked. For a negative control, 5 colonies harboring plasmid encoding for maltose-binding protein (MBP) were picked. After colony picking, toothpicks were removed, and plates were covered in foil and incubated at 37 °C for 14-16 hr in a ThermoFisher MaxQ 8000 shaker set to 250 rpm. Expression cultures (1 mL, TB-carb) were inoculated with 50 µL of starter cultures. In parallel, glycerol stocks were prepared in 350 µL 96-well plates (USA Scientific catalog #: 1830-9610) by mixing 50 µL of starter cultures with 50 µL sterile 1:1 glycerol:water. Glycerol stocks were sealed with cold-storage aluminum foil seals (VWR catalog #: 89049-034) and stored at -80 °C until hit identification. Inoculated expression cultures were incubated at 37 °C at 250 rpm for 3 hours. After 3 hours, plates were chilled on ice for 20 minutes before induction with IPTG

(0.5 mM final concentration), and then incubated at 18 °C at 250 rpm for 18-20 hours. Cells were pelleted by centrifugation (4000 rpm, 10 minutes, 4 °C). After supernatant was discarded, plates containing pelleted cells were sealed with silicone mats and stored at -20 °C for at least 24 hours before use.

### Library reactions in whole-cell

Frozen cell pellets in 96-well deep well plates were thawed on at room temperature for 10 minutes and then placed on ice. Cells were resuspended with 300 μL MES buffer (50 mM, pH 6.8). Substrate (125 mM in MES buffer, 96 μL, 20 mM final concentration), α-ketoglutarate disodium salt (250 mM in MES buffer, 96 μL, 40 mM final concentration), L-ascorbate (25 mM in water, 24 μL, 1 mM final concentration), and ferrous ammonium sulfate (12.5 mM, 24 μL, 0.5 mM final concentration) were added successively to resuspended cells. Plates were then covered loosely with foil and shaken for 24 hours at 25 °C. Reactions were quenched by addition of 300 μL acetonitrile. Plates were spun down to pellet cell debris and precipitated proteins. For individual well reaction analysis, 25 μL of the resultant supernatant was added to 225 μL of an 80:20 mixture of ACN:water with 10 mM ammonium acetate and 0.4% v/v ammonium hydroxide. Samples in later rounds of directed evolution were analyzed in a pooled manner. Pooled samples were prepared by combining 10 μL of six consecutive samples (A1-A6, A7-A12, etc) to a total volume of 60 μL. Each plate accounted for 16 pooled samples. The 60 μL sample mix was added too 190 μL of 80:20 mixture of ACN:water with 10 mM ammonium acetate and 0.4% v/v ammonium hydroxide. All LC/MS samples were filtered through a 96-well Pall AcroPrep™ Advance 96-well filter plate (350 μL, 0.2 μm Supor membrane), which was mounted onto a 96-well polypropylene sample plate (USA Scientific catalog #: 1830-9610). The stacked plates were spun down for 1 min at 4000 rpm. The plate containing filtrate was then sealed with heat-sealing aluminum foil. Samples were either stored at -20 °C or submitted directly to LC/MS analysis.

### Library analysis

LC/MS Analysis was carried out using a Waters Acquity 2D UHPLC coupled to a Waters Xevo-TQ mass spec. Column: Water BEH-Amide, 2.1x50mm 1.7 μm particle size equipped with a guard column. Mobile phase A: 100% $H_2O$ with 10 mM $NH_4CH_3COO^-$ and 0.04% $NH_4OH$. Mobile phase B: 95:5 ACN:water with 10 mM $NH_4CH_3COO^-$ and 0.04% $NH_4OH$.

Gradient:

| Time | A | B | Flow rate |
|---|---|---|---|
| Initial | 2.5 | 97.5 | 0.5 ml/min |
| 3 | 20 | 80 | -- |
| 4 | 2.5 | 97.5 | -- |
| 5.5 | 2.5 | 97.5 | -- |

Mass spec parameters – ESI mode: negative. Capillary voltage: 1 kV. Cone voltage: 30 V. Carboxylic acid product markers were directly injected into the source and the mass spec parameters were tuned for optimal ionization. The LC gradient conditions were optimized for separation of products from each other and from compounds in the reaction mixture. Multiple-reaction monitoring (MRM) was employed to detect specific fragments of product marker parent ions – specifically loss of water and formate. Ion counts were measured using Waters MassLynx.

### Library hit selection

For directed evolution rounds where every sample was analyzed by LC/MS, fold changes in ion counts for increased hydroxylation product were measured against the parent control samples. Mean turnover was calculated, and hits one standard deviation from the mean were selected for validation. For directed evolution rounds where samples were pooled, fold changes in ion counts for increased hydroxylation product were measured against a set of pooled parent samples. The mean fold change was calculated from parent, and pooled sets 1 SD from the mean were deconvoluted by analyzing individual samples. From the deconvoluted sample set, the same analysis relative to parent was carried out and hits were chosen if they met the 1 SD cutoff. In both cases, where samples were all analyzed individually or from pooled sets, typically ~5% of variants screened were selected for validation.

### Library hit validation

After selecting hits, cells from the corresponding glycerol stocks were streaked out on LB-carb agar plates. Streaked plates were incubated at 37 °C for 14-16 hours. Single colonies were picked for each hit to inoculate starter cultures (5 mL LB-carb). Starter cultures were grown at 37 °C for 10-12 hours. Plasmids were isolated from starter cultures and sent for sequencing. Unique variants were then re-expressed in 25-50 mL expressions. Variants were assayed using either clarified lysate or as pure enzymes isolated using the small-scale purification procedure described above. For validation with clarified lysates, enzyme concentration was estimated by SDS-PAGE using a calibration curve of protein at a known concentration followed by gel imaging on a LiCor Odyssey IR gel scanner and analysis of bands with ImageStudio Lite.

Small-scale validation reactions were carried out in 96-well deep well plates (Costar 3961). To enzyme (10-20 µM final concentration from purification or clarified lysate) in reaction buffer (MES 50 mM, pH 6.8) was added substrate (125 mM in MES buffer, 48 µL, 20 mM final concentration), α-ketoglutarate disodium salt (250 mM in MES buffer, 48 µL, 40 mM final concentration), L-ascorbate (25 mM in water, 12 µL, 1 mM final concentration), and ferrous ammonium sulfate (12.5 mM, 12 µL, 0.5 mM final concentration). Reaction plates were loosely covered with foil and incubated in a shaker at 25 °C for 24 hours. All reactions were run in triplicate. Reactions were quenched by addition of 150 µL acetonitrile. Total turnover (TTN) was calculated using a product marker calibration curve. In the case of product 4, calibration curves were made with both cis and trans product markers. Total turnover was calculated by dividing the product concentration by the enzyme concentration in the reaction. Stereoselectivity was calculated using the ratio of cis:trans TTNs. Samples for LC/MS analysis were prepared as described above. After LC/MS analysis, the variant with the highest mean fold change from parent that also retained 80% stereoselectivity was chosen as the winner of the round.

**Scale-up biocatalytic reaction with R2_11 H58F/L174G/V57H and product isolation**

To validate product identification, we performed a scale-up reaction, isolated the product, and characterized by NMR. To purified R2_11 H58F/L174G/V57H triple mutant (40 µM) in reaction buffer (MES 50 mM, pH 6.8) in a 50 mL Erlenmeyer flask was added cyclohexane carboxylic acid (Substrate **1**, 20 mM final concentration), α-ketoglutarate disodium salt (40 mM final concentration), L-ascorbate (1 mM final concentration), and ferrous ammonium sulfate (0.5 mM final concentration). The total reaction volume was 21 mL. The flask was shaken at 180 rpm at 25 °C for 24 hours. The reaction was quenched by acidification to pH ~1 with 1M HCl. The aqueous mixture was extracted with 2x15 mL of EtOAc. The combined organic layer was washed 3x15 mL with saturated brine, dried over sodium sulfate, and concentrated *in vacuo*. Purification by flash column chromatography afforded a 4-hydroxylcyclohexane carboxylic acid (**4**, 32 mg, 30% yield) as a mixture of *cis* and *trans* isomers, consistent with previous observations (Figure S4). $^1$H NMR (300 MHz, $D_2O$): δ3.79 (m, 1H), 3.52 (m, 1H), 2.41 (m, 1H), 2.29 (m, 1H), 1.89 (m, 4H), 1.74 (m, 2H), 1.57 (m, 6H), 1.35 (d, $J$ = 13 Hz, 2H), 1.18 (q, $J$ = 11 Hz, 2H) (Figure S6).

**Continuous fluorescence-based assay for Fe(II)/αKGs ("PBP assay") for initial rates measurements and Michaelis-Menten analyses**

**General set-up**

For Michaelis-Menten kinetic analysis of Fe(II)/αKGs used in this study, we developed a continuous coupled assay that takes advantage of a common mechanism in Fe(II)/αKGs.[5] Enzyme-catalyzed substrate oxidation is coupled to decomposition of α-ketoglutarate into succinate and $CO_2$. We used succinyl-CoA synthetase, which utilizes succinate, CoA, and ATP to generate succinyl-CoA along with ADP and inorganic phosphate ($P_i$). We detected $P_i$ by fluorescence emitted from an engineered phosphate binding protein (PBP, ThermoFisher Scientific PV4407).[6] Production of $P_i$ is equivalent to Fe(II)/αKG product generation (Figure S1). This assay can also be used to measure substrate uncoupled enzyme turnover. For comparison, a previously-reported Fe(II)/αKGs coupled assay used succinyl-CoA synthetase followed by two more coupled enzyme steps resulting in NADH consumption, which can be monitored with a continuous absorbance-based readout.[7]

Continuous fluorescence assays were carried out in 384 well-plates (Corning 3572) using a PerkinElmer EnVision plate reader. The top mirror module used was barcode 401. The excitation filter used was 405 nm (barcode 302) and the emission filter used was 450 nm (barcode 303). Typical assay components and amounts are shown in Table S2. Assays were set up in the following manner: a master mix was made fresh with substrate, buffer, $MgCl_2$, αKG, Fe(II), ascorbate, ATP, and Co-enzyme A. Master mix was distributed to individual wells, and PBP was added following addition of succinyl-CoA synthetase. This assay mixture was allowed to incubate at room-temperature for 5-10 minutes.

Commercial sources of αKG are contaminated with low levels of succinate, and this incubation period allows for consumption of succinate prior to addition of Fe(II)/αKG enzyme. Time-course assays were initiated by addition of Fe(II)/αKGs. For background negative controls, either no Fe(II)/αKG was added or no substrate was added. For the negative control where no substrate is added, any signal generated above the no Fe(II)/αKG control reactions is likely the result of substrate uncoupled Fe(II)/αKG turnover.[7] A calibration curve for the phosphate sensor was used to calculate [$P_i$] which is directly proportional to [product] from Fe(II)/αKG turnover.

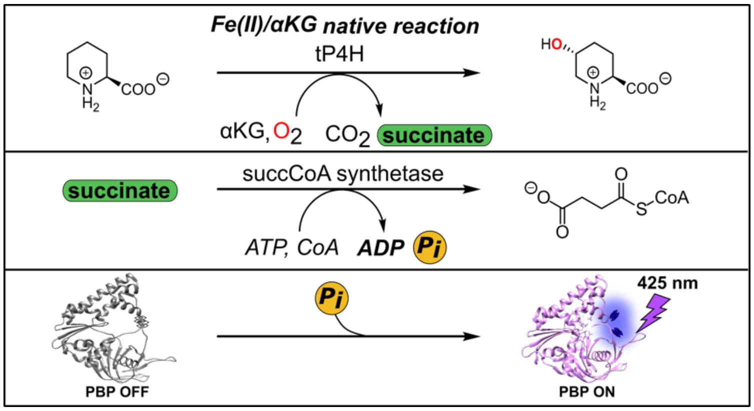

Figure S1. PBP assay general overview for Fe(II)/αKG oxidation reaction monitoring.

Table S2. General set-up for PBP assay on a 15 μL scale in a 384-well plate.

| Reagent | Stock concentration (mM) | Volume (μL) | Assay concentration | Unit |
|---|---|---|---|---|
| Substrate | 125 | 2.40 | 20 | mM |
| Asc | 25 | 0.30 | 0.5 | mM |
| Fe | 25 | 0.30 | 0.5 | mM |
| αKG | 10 | 0.4500 | 300.0 | μM |
| Fe(II)/αKG | 0.003 | 1.00 | 0.2 | μM |
| ATP (10 mM) | 10 | 0.45 | 0.3 | mM |
| CoA (10 mM) | 10 | 0.45 | 0.3 | mM |
| SucCD | 0.005 | 1.50 | 0.5000 | μM |
| $MgCl_2$ (100 mM) | 100 | 1.50 | 10.0 | mM |
| PBP (100 μM) | 0.1 | 3.00 | 20.000 | μM |
| Assay Buffer | 50 mM MOPS pH 7 50 mM NaCl | 3.65 | | |
| Total volume | | 15.00 | | |

### Reagents and sources

| Reagent | Source | Storage and composition |
|---|---|---|
| Substrate: Cyclohexane carboxylic acid | Sigma Aldrich Cat# 101834 | 125 mM in Assay Buffer, stored at r.t. |
| Substrate: L-pipecolic acid | TCI America Cat# P1404 | 125 mM in Assay Buffer, stored at r.t. |
| αKG | ChemImpex Cat# 00070 | 10 mM in assay buffer |
| L-ascorbate | Sigma Aldrich Cat# 255564-5G | Solid aliquots stored at r.t. – MilliQ water added at time of assay for 25 µM stock |
| ATP | ThermoFisher Scientific Cat# R1441 | 100 µL 10 mM stock diluted in MilliQ water, stored at -20 °C |
| Coenzyme A | Sigma Aldrich Cat# C3144 | Solid aliquots stored at -20 °C, cold Assay Buffer added at time of assay for 10 mM stock |
| Assay Buffer: 50 mM MOPS 150 mM NaCl | MOPS: MilliPore Sigma Cat# 475922100GM<br><br>NaCl: Fisher Scientific Cat# BP358-212 | Prepared using in-house MilliQ water, stored at r.t. |
| $MgCl_2$ | Fisher Scientific Cat# BP214 | 100 mM in MilliQ water, stored at r.t. |
| Fe(II) as ferrous ammonium sulfate | Sigma Aldrich Cat# 215406 | Solid aliquots stored at r.t., MilliQ water added at time of assay for 25 µM stock |
| Phosphate binding protein (PBP) | ThermoFisher Scientific Cat# PV4407 | Aliquoted and stored at -80 °C – diluted to 100 µM at time of assay in cold Assay Buffer |
| Succinyl CoA synthetase | Plasmid: Addgene cat# 83324; enzyme prepped in house | Aliquoted and stored in in 100 mM Tris, 150 mM NaCl, 20% glycerol pH 7.5 at -20 °C |

### Purification of succinyl-CoA synthetase

We found the lowest background fluorescence rate with succinyl-CoA synthetase prepared in-house after Ni-affinity purification followed by size exclusion chromatography (SEC). The SucCD gene was obtained from Addgene Cat# 83324. This plasmid did not encode for a 6xHis-tag fusion needed for Ni-affinity purification, so the SucCD gene was amplified and cloned into a standard pET22b vector between NdeI and XhoI restriction sites to add a C-terminal 6xHis-tag onto Chain D (α subunit) of SucCD.[8] SucCD was first Ni-affinity purified and dialyzed into 100 mM Tris, 150 mM NaCl, 20% glycerol pH 7.5. Ni-affinity purified SucCD was spin concentrated to to ~50-70 μM and loaded onto an AKTA FPLC equipped with an SEC column. In all preps, a higher MW species elutes from the column that are most likely SucCD oligomers based on SDS-PAGE analysis. Fractions containing purified heterotetrameric SucCD were identified by SDS-PAGE. Purified SucCD fractions were tested in the coupled assay to determine background noise. Any fractions containing higher MW species gave significant background fluorescence in the absence of Fe(II)/αKG enzyme. Therefore, only purified heterotetrameric SucCD was aliquoted and stored at -80 °C for assay use.

### Assay validation

We validated the fluorescence based coupled assay by comparing assay derived Michaelis-Menten parameters with literature reported values for GloF and HtyE. These results are shown in Table S3.

Table S3. Observed kinetic parameters for Fe(II)/αKGs HtyE and GloF compared to literature values.[9]

| Enzyme | Experimental $k_{cat}$ ($s^{-1}$) | Literature $k_{cat}$ ($s^{-1}$) | Experimental $K_M$ (mM L-Pro) | Literature $K_M$ (mM L-Pro) |
|---|---|---|---|---|
| GloF | 0.064 | 0.13 | 7.1 | 8.7 |
| HtyE | 0.18 | 0.65 | 8.3 | 4.2 |

### ProteinMPNN computational sequence redesign

#### ProteinMPNN multiple sequence alignment (MSA) input

To generate the MSA for tP4H redesign, four iterative HHblits[10] searches were performed against the UniRef30 database (accessed June 30, 2022 at E-value cutoffs of 1e-50, 1e-30, 1e-10, and 1e-4, and the final result was filtered for 90% identity redundancy, 50% coverage, and 30% minimum query identity.

#### Fixed Residue Selection

For tP4H, eight methods for fixed residue selection were employed, shown in Table S4. After testing five methods in the first set of experiments, three new methods (methods 6-8) were employed. Methods 1 and 2 fix residues near the active site. In methods 3-6, residues conserved in at least X% of sequences were determined by calculating the frequency of the native amino acid identity at each position in each multiple sequence alignment (MSA) sequence. If the amino acid identity was conserved in more than X% of sequences in the alignment, that position was fixed during design. Method 6 is a repeat of method 3 with two additional positions (228 and 230) near the binding pocket fixed. In methods 7 and 8, additional residues were fixed based on alternative conservation criteria. Highly conserved positions were determined by calculating the frequency of each amino acid at each position, identifying the most highly conserved amino acid, and ranking all positions by the frequency of the most highly conserved amino acid. The top 50% (method 7) or 70% (method 8) of positions were fixed during sequence design. For GriE, only method 8 was employed for design sequence generation after fixing active site residues. Complete lists of residues fixed in each of these methods are provided in the Supplementary spreadsheet (ProteinMPNN sequences_metrics tab).

Table S4. ProteinMPNN sequence generation methods for fixed residue selection.

| Method | Description |
|---|---|
| 1 | Fix active site residues:<br>tP4H: H58, Q92, K94, N96, K98, W106, H109, Q110, D111, F114, W115, Q173, H215, S217, R226, R247<br>GriE: Y59, Q93, K95, N97, K99, W107, H110, Q111, D112, V115, S116, V144, H169, V170, D171, P172, H210, A212, R221, L223, I225, R242, V246 |
| 2 | Fix active site (residues from Method 1) and 10 Å sphere around active site. The 10 Å sphere was defined as any residues containing sidechain atoms within 10 Å of any of the active site residues. |
| 3 | Fix active site (residues from Method 1) and residues conserved in at least 35% of MSA sequences |
| 4 | Fix active site (residues from Method 1) and residues conserved in at least 70% of MSA sequences |
| 5 | Fix active site (residues from Method 1) and residues conserved in at least 95% of MSA sequences |
| 6 | Method 3 + L228 and V230 |
| 7 | Fix active site (Residues shown in Method 1 + L228 and V230) and 50% most highly conserved residues from MSA |
| 8 | Fix active site (Residues shown in Method 1 + L228 and V230 for tP4H) and 70% most highly conserved residues from MSA.<br><br>For GriE redesign only this method was used. |

### ProteinMPNN Design of tP4H and GriE

The structure of tP4H was predicted with AlphaFold2[11] and used as structural input to ProteinMPNN.[12] Active site and conserved residues for tP4H and GriE were excluded from design as described in Table S4. Cysteine was excluded from the amino acid identities that could be installed during design. Three temperature sampling parameters (0.1, 0.2, and 0.3) were used during design.[13] A model of ProteinMPNN trained with 0.2 Å noise applied to training set protein backbones was used to perform sequence generation.

Sequences generated with ProteinMPNN were predicted with AlphaFold2, using model 3 with 6 recycling steps. Both designs and native tP4H and GriE predicted with low confidence if given only the single sequence and minimal recycling steps; we found that structural templating with MSAs was necessary for accurate prediction. To generate MSAs of each design for structure prediction, the MSA of the parent sequence was used, and the parent sequence was swapped for the design sequence. All sequences generated were predicted with Cα RMSD < 2.0 Å and pLDDT > 85.0 and were predicted to maintain critical structural features in the active site. For methods 6-8, we ordered the top ranked 32 out of 48 sequences based on top Cα RMSD values.

The following command was used to perform sequence design with ProteinMPNN on tP4H.

```
python $MPNN_PATH/protein_mpnn_run.py \
        --jsonl_path ../parsed_pdbs_bb.jsonl \
        --chain_id_jsonl ../assigned_chains.jsonl \
        --fixed_positions_jsonl ../masked_pos.jsonl \
        --out_folder $MPNN_OUTDIR \
        --num_seq_per_target 16 \
        --sampling_temp “0.1 0.2 0.3” \
        --batch_size 8 \
        --omit_AAs=’XC’
```

Where ../assigned_chains.jsonl contains the parsed PDB chain information: {“tP4H”: [[“A”]] }

This script generates 16 sequences for each sampling temp for a total of 48 sequences. The -omit_AAs line excludes cysteine residues from being installed during the design.

Sets of designs were distinguished by selection of fixed residues (Table S4).

### Design cloning and screening

All ProteinMPNN design sequences can be accessed in a supplementary Excel spreadsheet. Designs were purchased as IDT eBlocks and were used directly in In-Fusion reactions to construct design encoded plasmids. The eBlock 5’- and 3’-overhangs are shown below. eBlocks for tP4H were used directly in In-Fusion reactions with a pET-28a(+) vector that was digested with NcoI and XhoI. eBlocks for GriE were used directly in In-Fusion reactions with a pET-22b(-) vector that was digested with XbaI and XhoI. General eBlock designs are shown below:

General ProteinMPNN design eBlock DNA sequence and In-Fusion overhangs for pET-28a(+). 15 bp In-Fusion overhangs are bolded, and restriction sites are highlighted in red:

**5’ aggagatataccatg**ggcagcagccatcatcatcatcatcacagcagcggcctggttggcag c**DESIGNSEQUENCEHERE**c**tcgagcaccaccacc 3’**

General ProteinMPNN design eBlock DNA sequence and In-Fusion overhangs for pET-22b(-). 15 bp In-Fusion overhangs are bolded, and restriction sites are highlighted in red:

**5′ acaattcccctctagaaataattttgtttaactttaagaaggagatatacatDESIGNSEQUENCEHEREctcgagcaccaccacc 3′**

After In-Fusion reactions, constructed plasmids were transformed directly into BL21(DE3) cells. After a 1 hour outgrowth, starter cultures were prepared with 1 mL of LB in a 96-well deep well plate inoculated with transformed cells. Starter cultures were incubated in a shaker at 37 °C for 12-14 hours. Expression cultures (1 mL, TB-carb) were inoculated with 50 µL of starter cultures. In parallel, glycerol stocks were prepared in 350 µL 96-well plates (USA Scientific catalog #: 1830-9610) by mixing 50 µL of starter cultures with 50 µL sterile 1:1 glycerol:water. Glycerol stocks were sealed with cold-storage aluminum foil seals (VWR catalog #: 89049-034) and stored at -80 °C until design hit identification. Glycerol stocks were sealed with cold-storage aluminum foil seals (VWR catalog #: 89049-034) and stored at -80 °C until hit identification. Inoculated expression cultures were incubated at 37 °C at 250 rpm for 3 hours. After 3 hours, plates were chilled on ice for 20 minutes before induction with IPTG (0.5 mM final concentration), and then incubated at 18 °C at 250 rpm for 18-20 hours. Cells were pelleted by centrifugation (4000 rpm, 10 minutes, 4 °C). After supernatant was discarded, plates containing pelleted cells were sealed with silicone mats and stored at -20 °C for at least 24 hours before use. Expression of designs was tested with SDS-PAGE. The general workflow for assessing hits for tP4H ProteinMPNN (Methods 6-8, Table S4) is shown in Figure S2. A similar workflow was used for GriE. Designs passing each criteria can be found in Supplementary Spreadsheet – ProteinMPNN sequences_metrics.

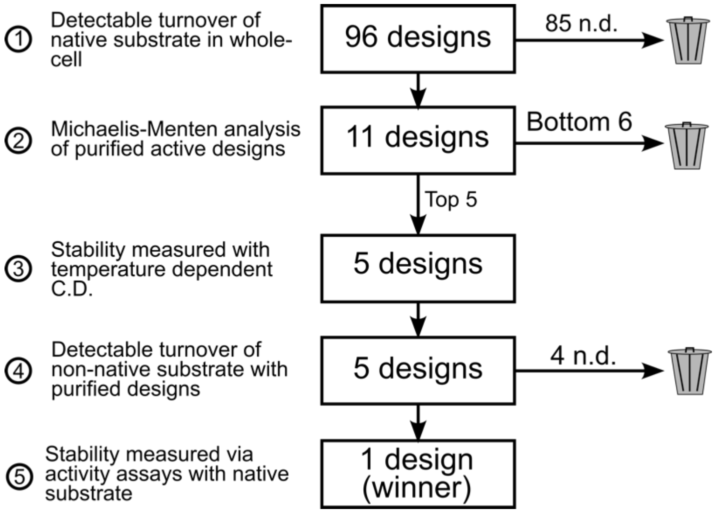

Figure S2. Selection criteria and workflow for the second set of tP4H ProteinMPNN design screening.

**Initial activity screen in whole-cell:** Initial screening was carried out to identify designs that turned over the native tP4H and GriE substrates L-pipecolic acid and L-leucine, respectively. Frozen cell pellets in 96-well deep well plates were thawed on at room temperature for 10 minutes and then placed on ice. Cells were resuspended with 300 µL MES buffer (50 mM, pH 6.8). Amino acid substrate (125 mM in MES buffer, 96 µL, 20 mM final concentration), α-ketoglutarate disodium salt (250 mM in MES buffer, 96 µL, 40 mM final concentration), L-ascorbate (25 mM in water, 24 µL, 1 mM final concentration), and ferrous ammonium sulfate (12.5 mM, 24 µL, 0.5 mM final concentration) were added successively to resuspended cells. Plates were then covered loosely with foil and shaken for 24 hours at 25 °C. Reactions were quenched by addition of 300 µL acetonitrile. Plates were spun down to pellet cell debris and precipitated proteins. For individual well reaction analysis, 25 µL of the resultant supernatant was added to 225 µL of an 80:20 mixture of ACN:water with 10 mM ammonium acetate and 0.4% v/v ammonium hydroxide. LC/MS was run using the method described above in the Site Saturation Mutagenesis Library Analysis workflow.

**Michaelis-Menten kinetics screen with purified tP4H designs:** Any designs that turned over native substrates were expressed and purified using the small-scale Ni-affinity purification protocol described above. Initial rates were measured using the PBP assay described above. For tP4H, Michaelis-Menten analysis was carried out where the $K_M$ for each design for αKG was determined first by fixing L-pipecolic acid at 20 mM and measuring initial rates with the following concentrations of αKG: 0, 6, 19, 56, 167 and 500 μM. For Michaelis-Menten analysis with tP4H using L-pipecolic acid, a saturating concentration of αKG was used along with 200 nM enzyme. Initial rates were then measured at the following concentrations of L-pipecolic acid: 0, 0.25, 0.74, 2.2, 6.7, 20, and 40 mM. The top 5 designs with the highest $k_{cat}$ were selected for analysis by temperature dependent circular dichroism (CD) spectroscopy (Figure S8).

**Michaelis-Menten kinetics with purified GriE designs:** Any GriE designs that turned over L-leucine with yield at least 2-fold below the wild-type GriE control were selected for purification and initial rate analysis. This criteria included 27 enzymes out of 32 designs. At this stage only observed rates were measured at 300 μM aKG and 20 mM L-leucine, without varying concentrations, to determine Michaelis-Menten parameters. After the initial rates were measured, the top 5 were selected for analysis by temperature dependent circular dichroism (CD) spectroscopy. After temperature dependent CD, the most stable and active design GM_A9 was characterized with a full Michaelis-Menten analysis. The $K_M$ of αKG for GM_A9 was determined first by fixing L-leucine at 20 mM and the following αKG concentrations were used with 200 nM enzyme to measure initial rates: 800, 400, 200, 100, 50, 25, and 12.5 μM. Then, a saturating concentration of 300 μM αKG was used along with 200 nM enzyme. Initial rates were measured at the following concentrations of L-leucine: 10, 3.3, 1,1, 0.37, 0.12, 0.04, and 0.01 mM.

**Temperature dependent CD spectroscopy:** To determine secondary structure and thermostability of design candidates from the second-pass screen described above, CD measurements were carried out on a JASCO J-1500 instrument using a 1 mm pathlength cuvette. Samples of purified protein were prepared at 0.4 mg/mL in 50 mM potassium phosphate buffer, pH 7.0. The sample temperature was ramped from 25 °C to 95 °C with full spectrum scans from 190 nm to 260 nm performed after each 10 °C interval. The molar residue ellipticity (MRE) at 220 nm was plotted over the temperature gradient to visualize temperature of unfolding. Representative CD data for tP4H and GriE can be found in Figure S8 and Figure S12B, respectively.

**Non-native activity screen with purified enzyme:** To determine if designs reacted with a non-native free acid substrate, purified enzymes were tested in small-scale biocatalytic reactions with cyclohexane carboxylic acid (substrate **1**). Reactions were run in the same manner described in SSM Library Hit Validation. LC/MS was used to determine TTN values.

**Stability-activity assay with purified enzyme:** Design stability was assessed by measuring initial rates at 25, 45 and 65 °C at t=0, 7 hours, and 24 hours unless otherwise stated. Initial rates were measured using the PBP assay described above.

**SDS-PAGE of tP4H variants and ProteinMPNN redesigns**

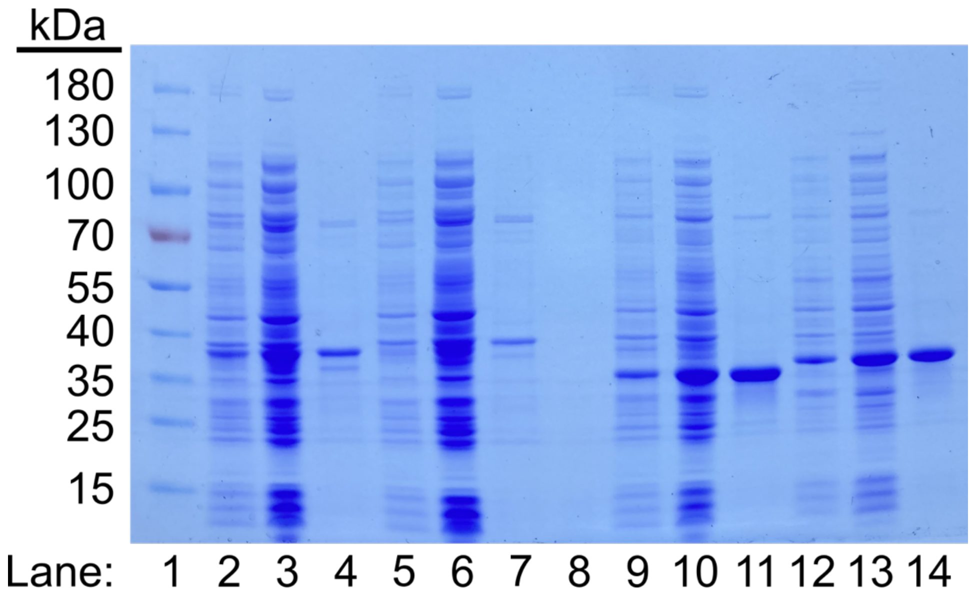

Figure S3. SDS-PAGE gel of wild-type tP4H, tP4H H58L/W170Q/E118K, R2_11, and R2_11 H58F/L174G/V57H. For each construct, gel samples are shown for post-expression from cell culture, lysate, and after Ni-affinity purification. All samples collected were diluted in the same way, so differences in staining intensity reflect differences in sample protein concentration.

Table S5. Gel lanes and descriptors for Figure S3.

| Lane | Description | Lane | Description |
|---|---|---|---|
| 1 | Ladder | 8 | Blank |
| 2 | wt tP4H post expression | 9 | R2_11 base design post expression |
| 3 | wt tP4H lysate | 10 | R2_11 base design lysate |
| 4 | wt tP4H purified | 11 | R2_11 base design purified |
| 5 | tP4H H58L/W170Q/E118K post expression | 12 | R2_11 H58F/L174G/V57H post expression |
| 6 | tP4H H58L/W170Q/E118K lysate | 13 | R2_11 H58F/L174G/V57H lysate |
| 7 | tP4H H58L/W170Q/E118K purified | 14 | R2_11 H58F/L174G/V57H purified |

### LC-MS characterization of substrate 1 hydroxylation products and calibration curves

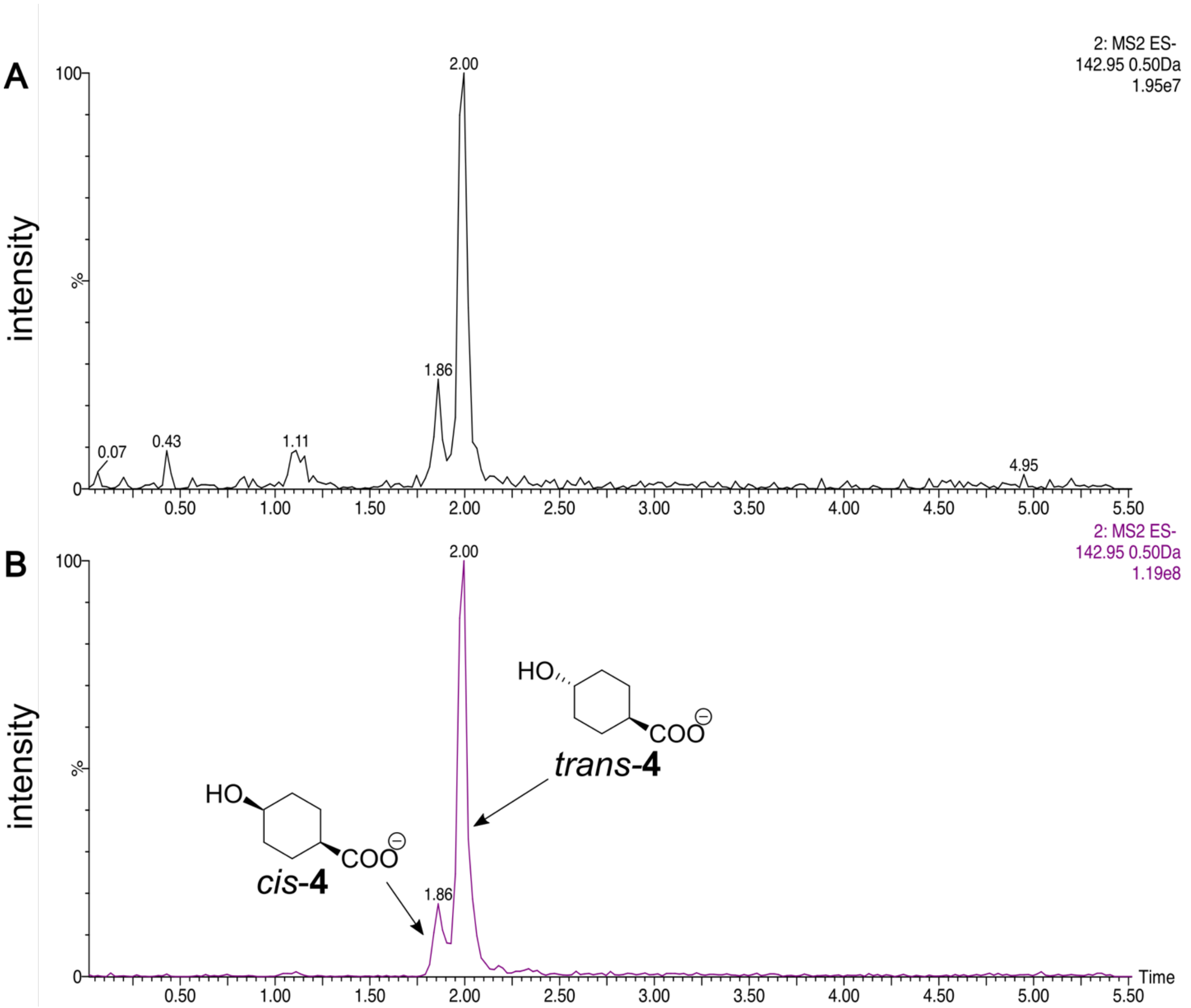

Figure S4. A) LC-MS extracted ion chromatogram (EIC) of a representative reaction sample with the R2_11 H58F/L174G/V57H triple mutant using the method described above in "Library Analysis." The 4:1 ratio of *trans* to *cis* was determined from calibration curves constructed from co-injected markers at varying ratios (Figure S5). The 4:1 selectivity was consistent across all mutants from both wild-type tP4H and R2_11. B) LC-MS EIC of a co-injection of authentic *cis*-**4** and *trans*-**4** product markers using the same method

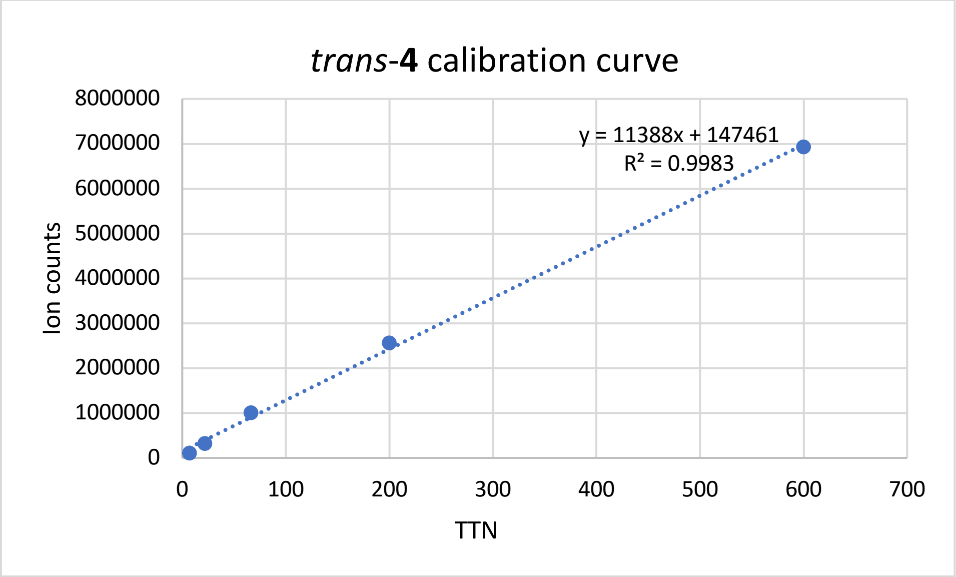

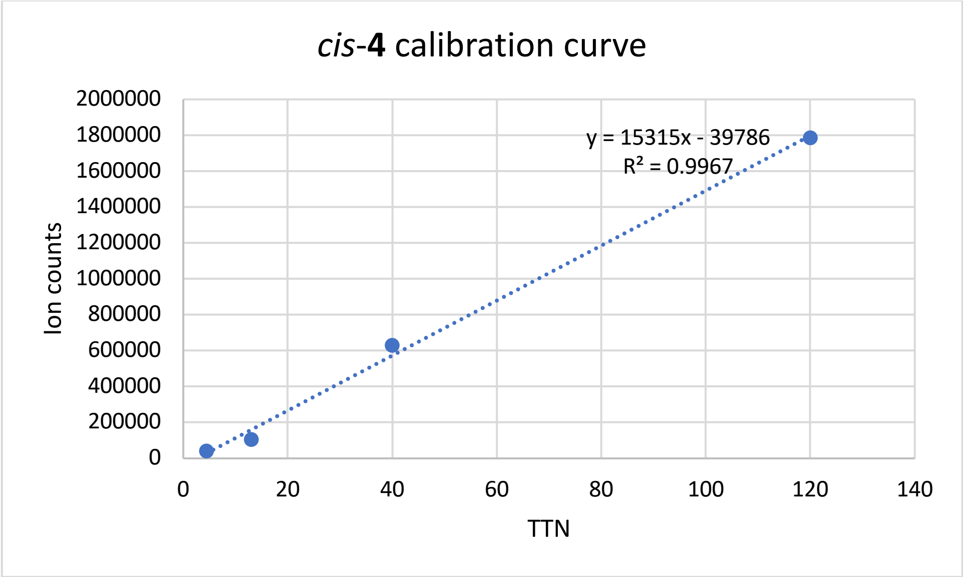

Figure S5. Calibration curves for authentic trans-4-hydroxycyclopentane carboxylic acid product markers. Ion counts were obtained from the extracted ion chromatograms for m/s 142.95 ([M]-1) using the LC-MS method described in the "Library Analysis" section above. A) Calibration curve using product marker for *trans*-4-hydroxycyclohexane carboxylic acid (Combi-blocks Cat. #: OR-5210). B) Calibration curve using product marker for *cis*-4-hydroxycyclohexane carboxylic acid (Combi-blocks Cat. #: QG-7784).

### NMR spectra of isolated product 4 and product markers

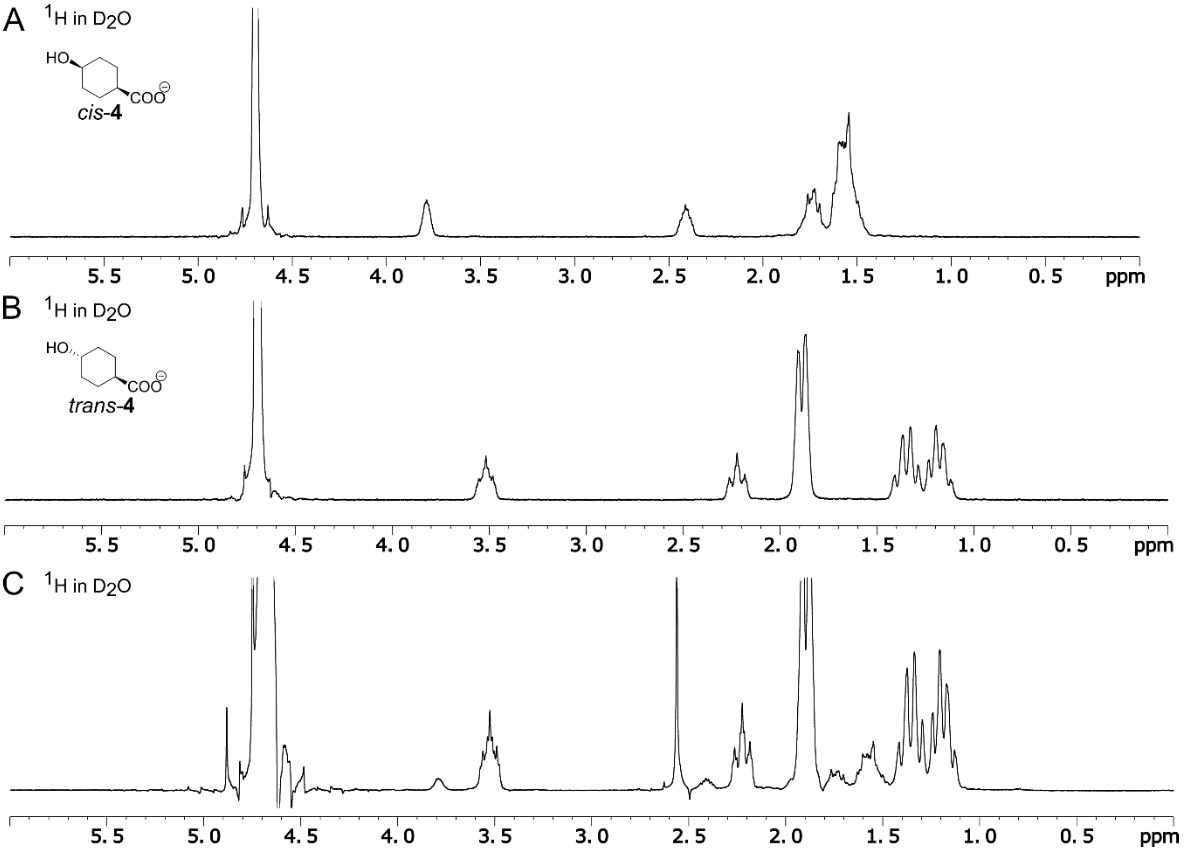

Figure S6. A) $^{1}H$ NMR spectrum (300 MHz, $D_2O$) of *cis*-**4** authentic product marker (Combi-blocks Cat. #: QG-7784). B) $^{1}H$ NMR spectrum (300 MHz, $D_2O$) of *trans*-**4** authentic product marker (Combi-blocks Cat. #: OR-5210). C) $^{1}H$ NMR spectrum (300 MHz, $D_2O$) of isolated **4** as a mixture of *cis* and *trans* isomers after flash silica gel chromatography from a scale-up biocatalytic reaction with R2_11 H58F/L174G/V57H triple mutant.

### LCMS traces for reactions of GriE and GM_A9

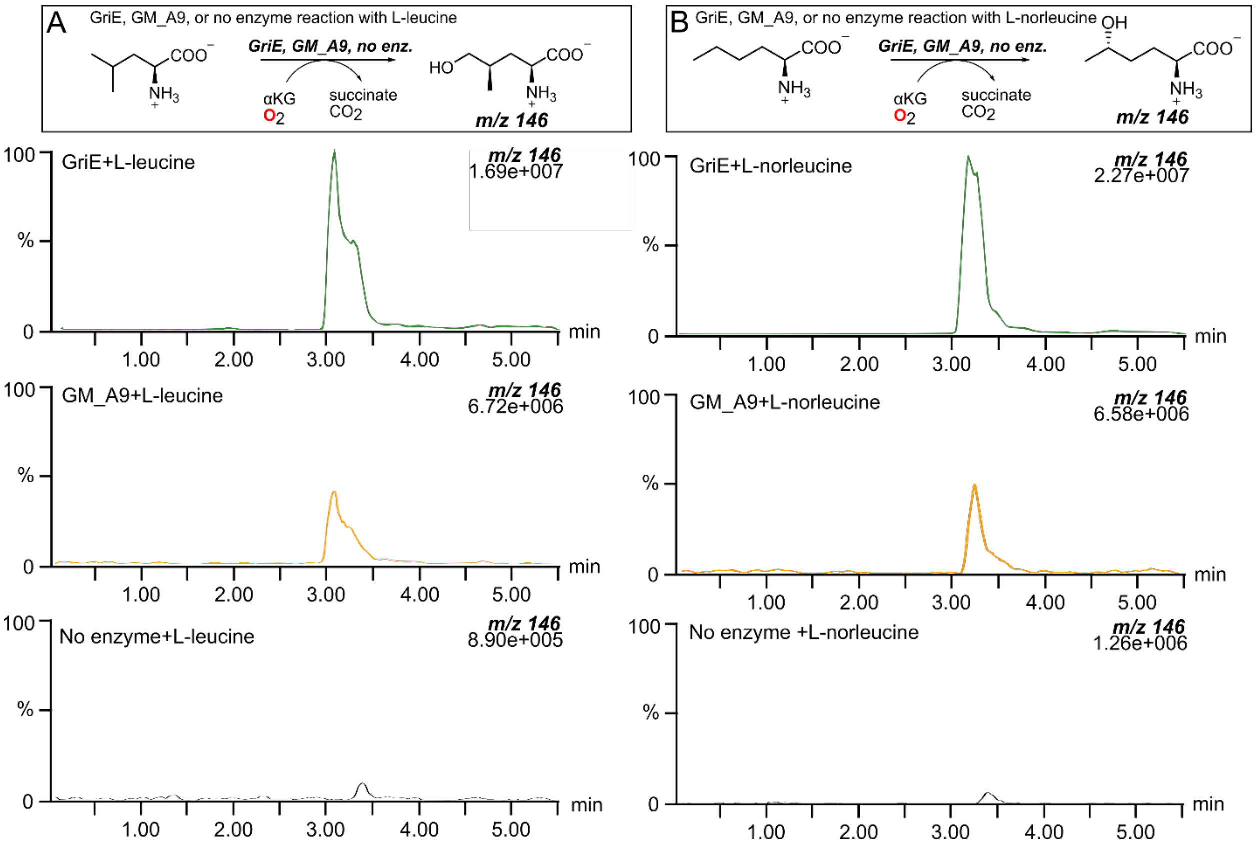

Figure S7. LCMS data were collected on a Waters Xevo TQ using the method described under "Library Analysis." Reactions were carried out with purified enzyme (15 μM) in reaction buffer (MES 50 mM, pH 6.8) with substrate (20 mM), αKG (40 mM), ferrous ammonium sulfate (0.5 mM) and ascorbic acid (1 mM). The extracted m/z as well as ion intensity are listed in the upper right-hand corner of each trace. A) Extracted ion chromatograms for reaction samples of GriE with L-leucine, GM_A9 with L-leucine, and a no enzyme with L-leucine control. B) Extracted ion chromatograms for reaction samples of GriE with L-norleucine, GM_A9 with L-norleucine, and a no enzyme with L-norleucine control.

### Circular dichroism data for wild-type tP4H and redesigned variants

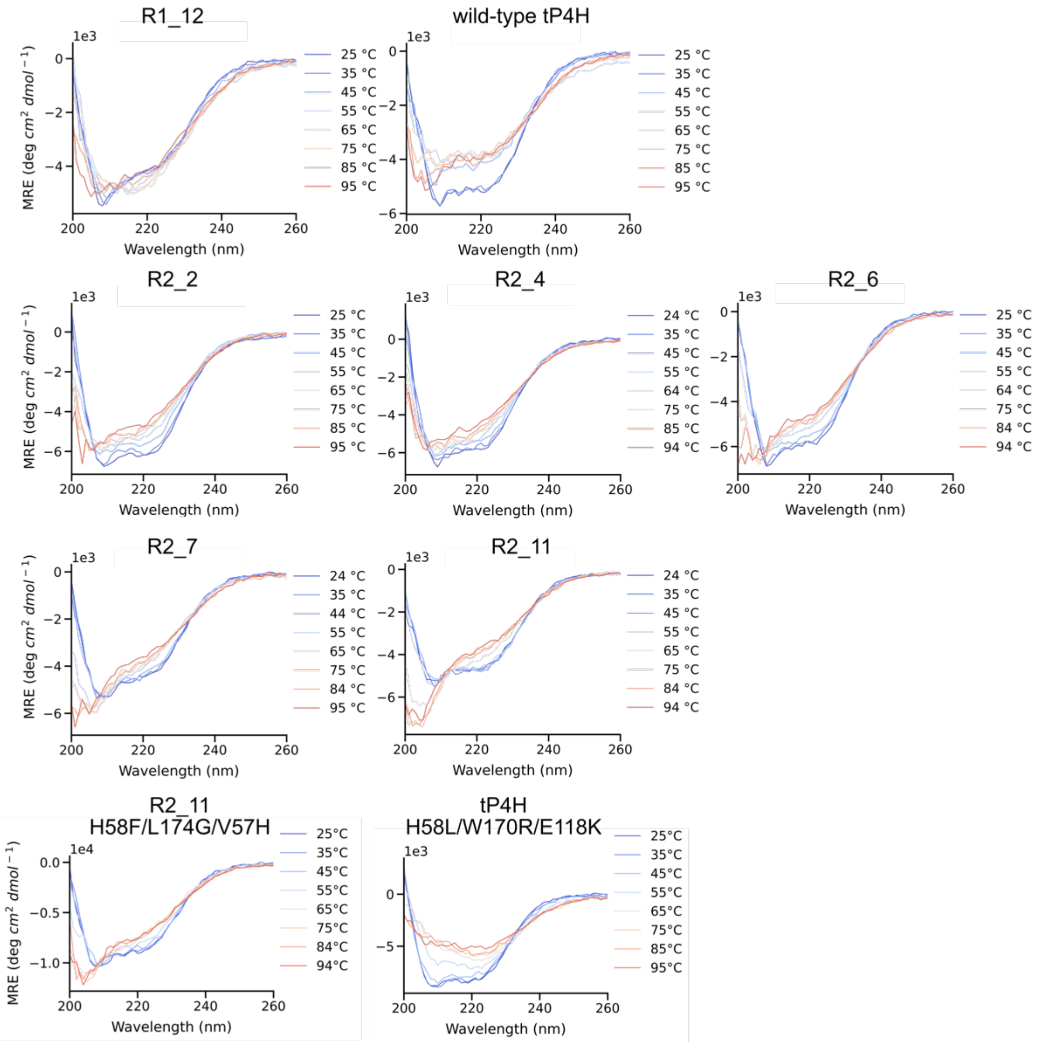

Figure S8. CD spectra for wild-type tP4H and redesigns, as well as tP4H and R2_11 triple mutants. CD spectra were collected at 25-95°C in 10 °C increments. Protein was prepared at 0.4 mg/mL in potassium phosphate buffer (50 mM, pH 7.0).

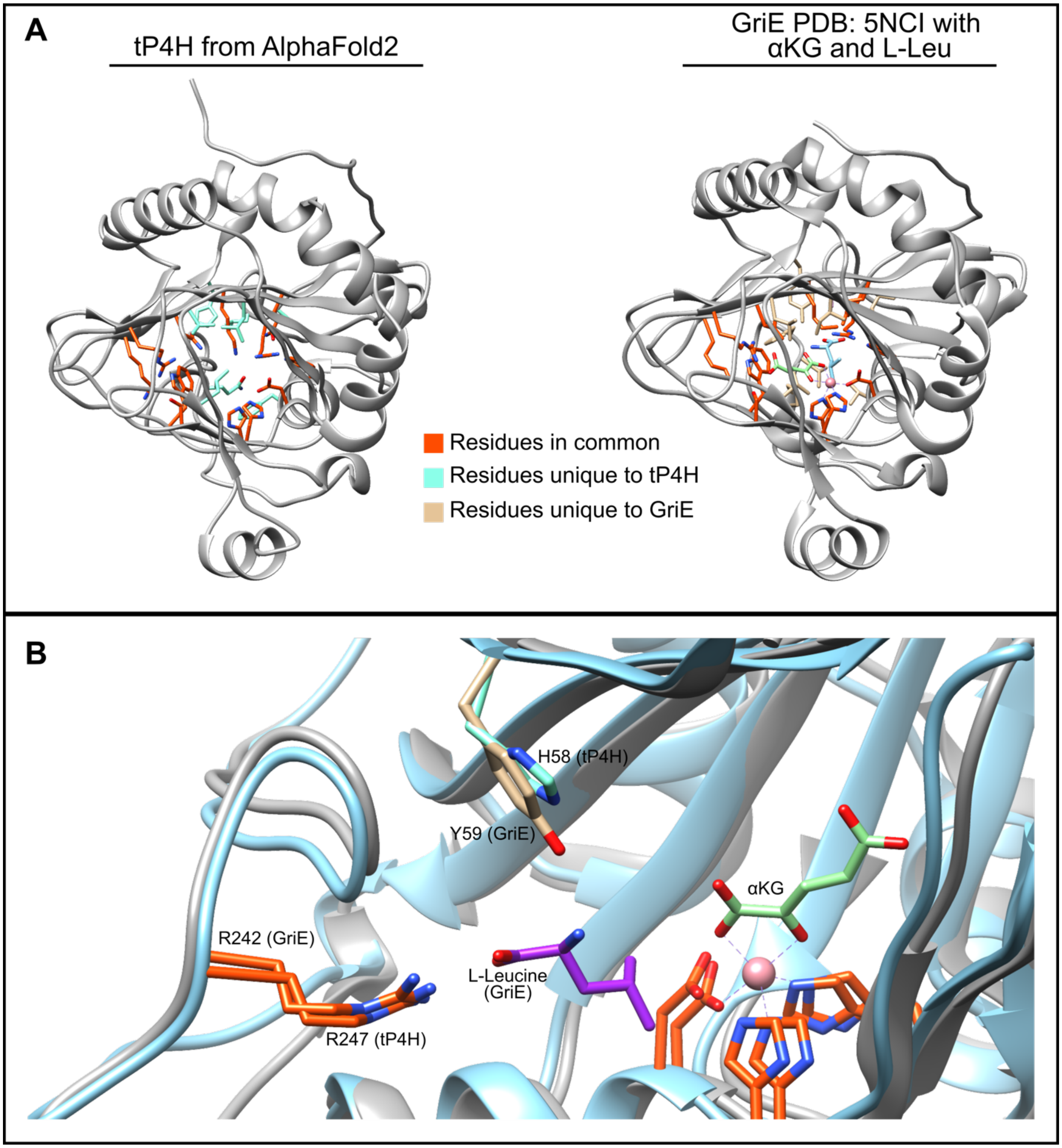

Figure S9. A) Structural model of tP4H from AlphaFold2 compared to a crystal structure of Fe(II)/αKG GriE (PDB ID: 5NCI).[3] Both common and unique features are shown via colored side-chains. The GriE structure contains bound cobalt (metal that GriE was crystallized with, light pink), αKG (light green), and the L-Leucine substrate (purple). B) Active site overlay of GriE (grey) and tP4H (light blue). If the L-pipecolic acid substrate in tP4H binds similarly to the orientation of L-Leucine in GriE, then the carboxylate group would likely interact with the tP4H R247 residue, and the amine group would likely interact with tP4H H58.

Table S6. Wild-type Fe(II)/αKG enzyme turnover with native substrates in whole-cell reactions.

| Fe(II)/αKG | Substrate | % conversion to hydroxylated product[a,b] |
|---|---|---|
| PolL | L-leucine | 59 |
| LdoA | L-leucine | <5 |
| IDO | L-leucine | 92 |
| GriE | L-leucine | 84 |
| Gox | L-leucine | 93 |
| cP3H | L-proline | 45 |
| cP4H | L-proline | 19 |
| GloF | L-proline | 59 |
| HtyE | L-proline | 68 |
| tP4H[c] | L-pipecolic acid | 17 |
| GetF | L-pipecolic acid | >99 |
| PiFa | L-pipecolic acid | 52 |

[a] All reactions were carried out in whole-cell with 20 mM substrate, 40 mM αKG, 1 mM ferrous ammonium sulfate and 1 mM ascorbic acid for 24 hours at 25 °C. Frozen and thawed cell pellets were resuspended in buffer (MOPS 50 mM pH 7.0) at 5% of the expression volume.
[b] Conversion to hydroxylated amino acid products was quantified by analytical HPLC-UV analysis after Fmoc-Cl derivatization of reaction mixtures.
[c] Previous reports with tP4H suggest that the enzyme needs to be co-expressed with a chaperone in order to have activity.[1] We found that chaperone co-expression was not necessary and enzyme activity with and without chaperone co-expression was comparable. Additionally, we found no difference in the initial rates of tP4H with L-pipecolic acid with and without chaperone co-expression.

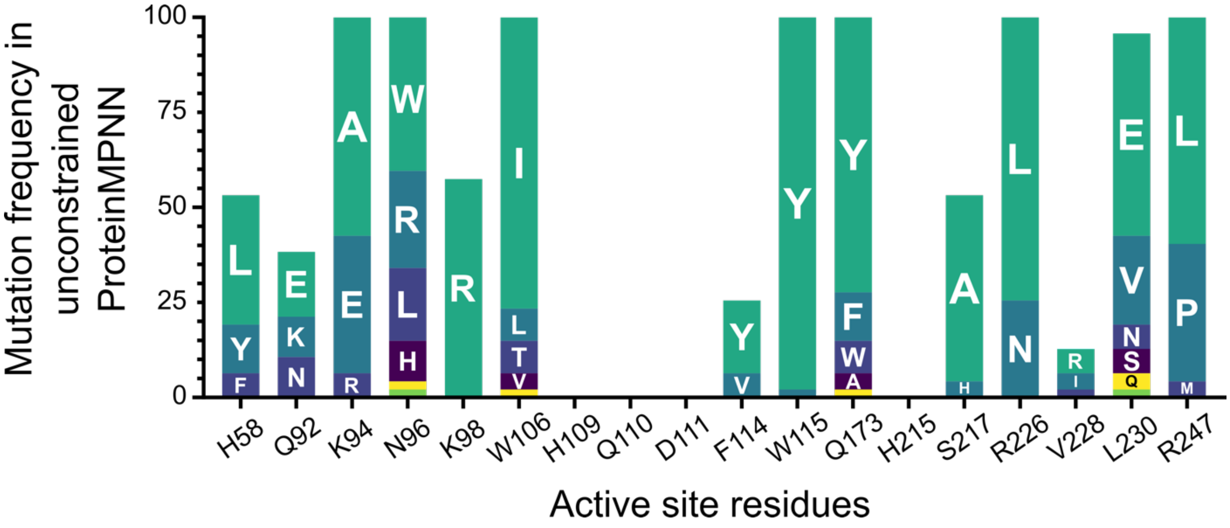

Figure S10. Unconstrained ProteinMPNN redesigns tP4H active site residues. The x-axis shows key tP4H active site residues that were fixed in typical tP4H ProteinMPNN redesigns (see Table S4). The y-axis shows the frequency of designed mutations at these sites when ProteinMPNN is unconstrained (i.e. no active site residues fixed). Specifically, the colored bars indicate the frequency across 48 output sequences for any amino acid other than the wild-type parent. We observed frequent mutations at functionally important sites. For example, position 94 is critical for αKG cofactor binding and is changed from the wild-type residue K in 100% of the unconstrained ProteinMPNN redesigns. Notably, unconstrained ProteinMPNN did not introduce mutations at $Fe^{2+}$ ligands H109, D111, and H215 (Figure 2) or the nearby residue Q110. ProteinMPNN receives only protein information as input and does not explicitly account for the presence of the metal ion,[12] so this motif may be preserved due to its frequent presence in the original training set.

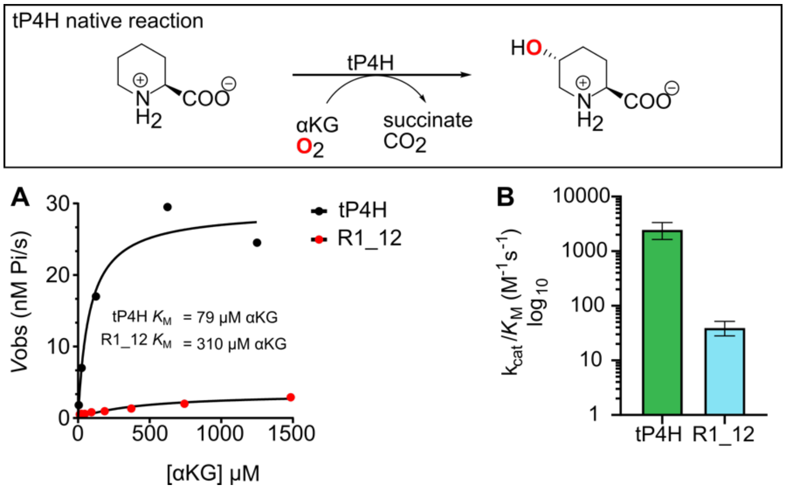

Figure S11. A) Michaelis-Menten analysis of wild-type tP4H and R1_12 for αKG using the native reaction with L-pipecolic acid with 200 nM enzyme. B) Catalytic efficiency comparison of wild-type tP4H and ProteinMPNN design candidate R1_12. Initial rates were measured using the PBP assay. Initial rates were plotted against substrate concentrations in GraphPad Prism and the non-linear Michaelis-Menten fit function was used to calculate $k_{cat}$ and $K_M$ values. To calculate the standard error of the fit for $k_{cat}/K_M$, an alternative form of the Michaelis-Menten equation was used: $V_{obs}=(k_{cat}/K_M)[E]_0[S]/(1+([S]/K_M))$.

Table S7. Michaelis-Menten parameters for active tP4H ProteinMPNN designs[a]

| | wt | R2_1 | R2_2 | R2_3 | R2_4 | R2_5 | R2_6 | R2_7 | R2_8 | R2_9 | R2_11 |
|---|---|---|---|---|---|---|---|---|---|---|---|
| $k_{cat}$ ($s^{-1}$) | 0.17 | 0.027 | 0.041 | 0.005 | 0.068 | 0.037 | 0.06 | 0.037 | 0.016 | 0.033 | 0.16 |
| $K_M$ (mM) | 0.5 | 41 | 1.3 | <0.25 | 2.1 | 51 | 3.7 | 1.4 | 2.6 | 1 | 6.7 |
| $k_{cat}/K_M$ ($M^{-1}*s^{-1}$) | 330 | 0.7 | 31 | -- | 32 | 0.7 | 16 | 27 | 6.2 | 32 | 24 |

[a] Active tP4H ProteinMPNN designs obtained from Methods 6-8 (Table S4). Initial rates (product vs. time curves) were measured using the PBP assay using 200 nM enzyme. First-derivatives of product vs time curves were plotted against substrate concentrations in GraphPad Prism and the non-linear Michaelis-Menten fit function was used to calculate $k_{cat}$ and $K_M$ values. For R2_3, observed rates exhibited saturation behavior for the lowest substrate concentration tested (0.25 mM). For this variant only a limit for $K_M$ could be determined and $k_{cat}/K_M$ could not be measured.

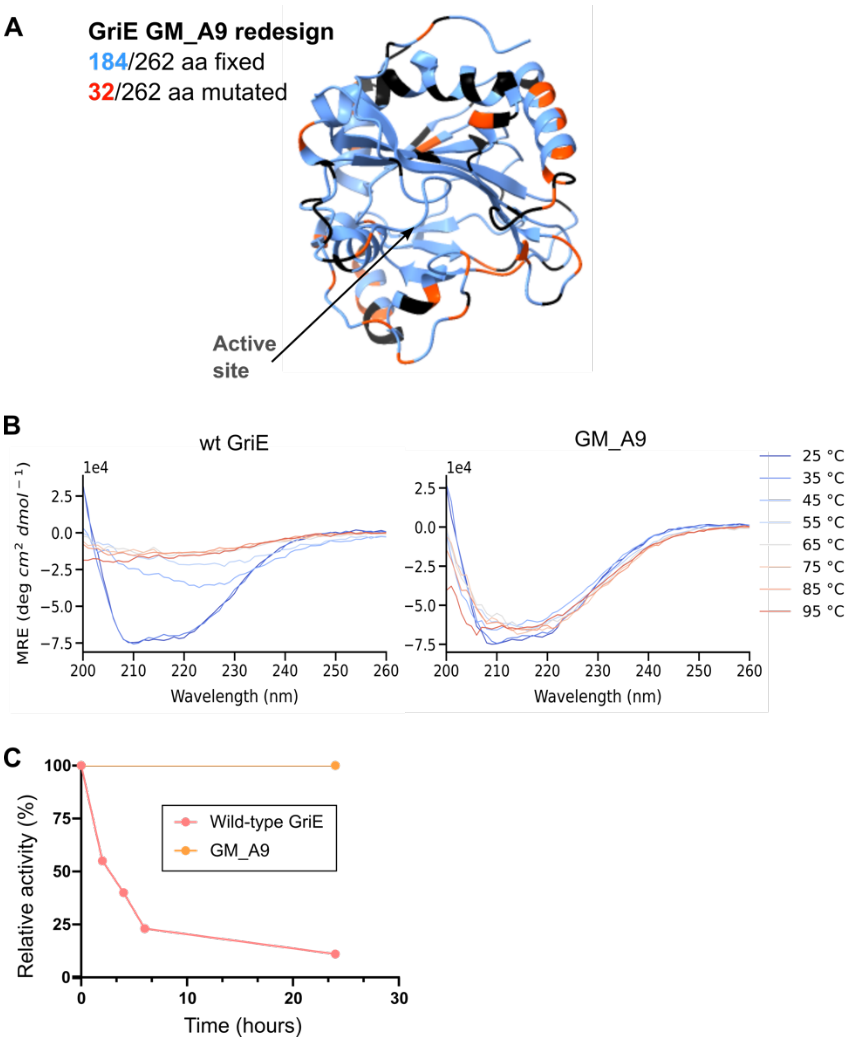

Figure S12. A) GriE structure (PDB 5NCI)[3] color-coded to show sites fixed in the design process (blue, Supplementary spreadsheet – ProteinMPNN sequences_metrics) and sites mutated in the ProteinMPNN GM_A9 redesign (orange-red). Sites colored black were neither fixed nor redesigned in the GM_A9 variant. B) Temperature dependent CD of wild-type GriE compared to ProteinMPNN redesign GM_A9. C) Activity-stability analysis of wild-type GriE compared to ProteinMPNN redesign GM_A9 at room temperature. Activity measurements were made using the PBP coupled assay described above in the Methods section.

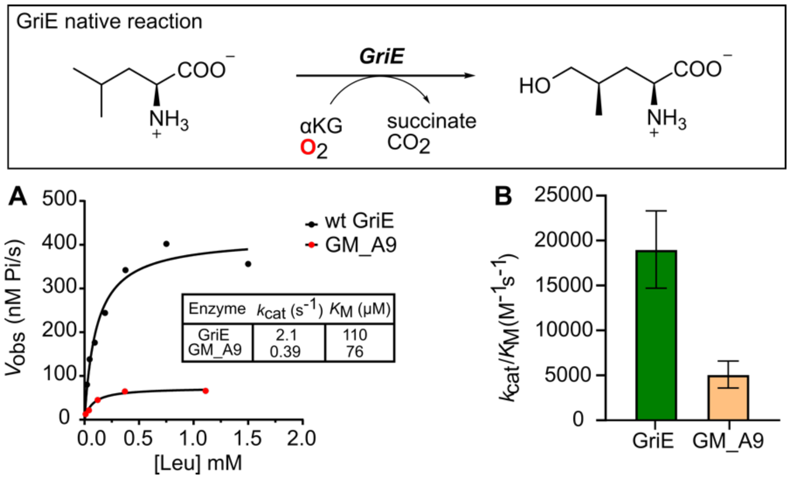

Figure S13. (A) Michaelis-Menten analysis of wild-type GriE and ProteinMPNN redesign GM_A9 with L-Leucine with 200 nM enzyme. (B) Catalytic efficiency comparison of wild-type GriE and ProteinMPNN redesign GM_A9. Error bars report the standard error of the fit calculated from 95% confidence interval values for $k_{cat}$ and $K_M$. Initial rates were measured using the PBP assay described aboce under Materials and Methods. Initial rates were plotted against substrate concentrations in GraphPad Prism and the non-linear Michaelis-Menten fit function was used to calculate $k_{cat}$ and $K_M$ values. To calculate the standard error of the fit for $k_{cat}/K_M$, an alternative form of the Michaelis-Menten equation was used: $V_{obs}=(k_{cat}/K_M)[E]_0[S]/(1+([S]/K_M))$.

Products[a]

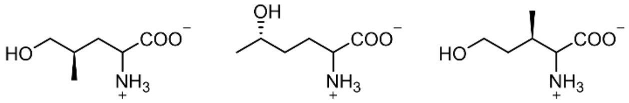

From substrates: L-leucine L-norleucine L-allo-isoleucine

| Enzyme | TTN values | | |
|---|---|---|---|
| GriE[b] | 7800 | 1800 | 190 |
| GM_A9[c] | 700 | 420 | n.d. |
| Fold-decrease rel. to wild-type | 11x | 4.3x | NA |

Table S8. Substrate scope of ProteinMPNN redesign GM_A9 compared to wild-type GriE.

[a] Products were determined by comparing the retention times of M+16 products of GM_A9 with wild-type GriE samples.

[b] TTN values for wild-type GriE were taken from a previous publication.[2]

[c] TTN values for GM_A9 products were determined by taking the M+16 product area% value compared to substrate.

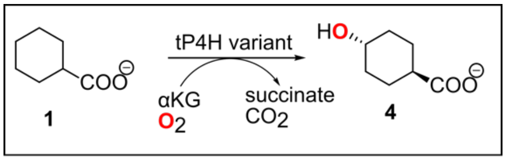

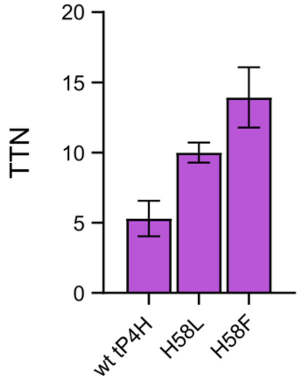

Figure S14. TTN effects for H58L and H58F mutations in wild-type tP4H. H58L is the round 1 winner for directed evolution from wild-type tP4H. H58F is the round 1 winner for directed evolution from the R2_11 variant. Reactions were carried out for 24 hrs at 25 °C using purified enzyme (20 µM) in MES buffer (50 mM, pH 6.8), with 20 mM cyclohexane carboxylic acid **1**, 40 mM αKG, 1 mM ferrous ammonium sulfate, and 1 mM ascorbic acid. Concentration of **4** in quenched reaction samples was quantified by analytical LC-MS analysis.

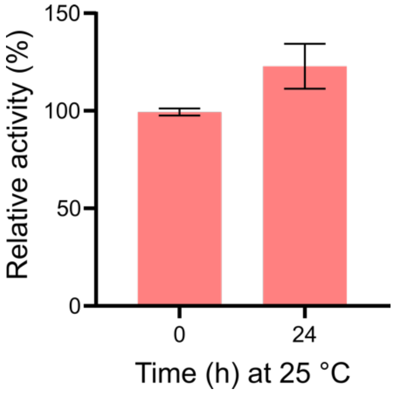

Figure S15. Activity-stability analysis of R2_11 H58F/L174G/V57H at room temperature. Relative activity was determined using PBP assay described above.
